# Time-dependent thermodynamic relationships for a Brownian particle that walks in a complex network

**DOI:** 10.1101/2023.12.06.570486

**Authors:** Mesfin Asfaw Taye

## Abstract

The thermodynamics feature of systems that are driven out of equilibrium is explored for *M* Brownian ratchets that are arranged in a complex network. The exact time-dependent solution depicts that as the network size increases, the entropy *S*, entropy production *e*_*p*_(*t*), and entropy extraction *h*_*d*_(*t*) of the system step up which is feasible since these thermodynamic quantities exhibit an extensive property. In other words, as the number of lattice size increases, the entropy *S*, entropy production *e*_*p*_(*t*), and entropy extraction *h*_*d*_(*t*) step up revealing that these complex networks can not be reduced into the corresponding one-dimensional lattice. On the contrary, the rate for thermodynamic relations such as the velocity V, entropy production rate *ė*_*p*_(*t*) and entropy extraction rate 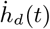 become independent of the network size in the long time limit. The exact analytic result also shows that the free energy decreases with the system size. The model system is further analyzed by including heat transfer via kinetic energy. Since the heat exchange via kinetic energy does not affect the energy extraction rate, the heat dumped to the cold reservoirs contributes only to the internal entropy production. As the result, such systems exhibit a higher degree of irreversibility. The thermodynamic features of a system that operates between hot and cold baths are also compared and contrasted with a system that operates in a heat bath where its temperature varies linearly along the reaction coordinate. Regardless of the network arrangements, the entropy, entropy production, and extraction rates are considerably larger for the linearly varying temperature case than a system that operates between hot and cold baths.

**PACS numbers:** Valid PACS appear here

## I. INTRODUCTION

Thermodynamics is an important discipline that historically began to study how heat and matter exchange from one system to another system. Since its relevance to science and technology is considerably high, the study of thermodynamics systems have obtained considerable attention. Particularly, exploring the dependence of entropy production *e*_*p*_ is vital since not only it signifies the degree of irreversibility of a given system but also serves as a fundamental limit regarding the efficiency of thermodynamic systems. As indicated by the second law of thermodynamics, *e*_*p*_ ≥ 0 signifies that systems always maximize their entropy production during an irreversible process and *e*_*p*_ = 0 for reversible processes [1, 2]. In the past decades, several frameworks have been developed to tackle how entropy production, as well as entropy production rates behave. Some pioneering work in this regard includes Onsager’s reciprocity theory [3, 4] which applies to a system operating near a linear response regime.

Systems that are genuinely far from equilibrium also have obtained considerable attention see for example the works [5–21]. Some of these studies have explored the dependence of the thermodynamic quantities either by considering classical systems [22, 23] or systems that operate in the quantum realm [24–26]. Since the Boltzmann-Gibbs nonequilibrium entropy along with the entropy balance equation still remain valid for systems that are far from equilibrium, exploiting this property, many theoretical studies have been conducted to study the entropy production in terms of the local probability density and transition probability rate. For discrete systems, employing the master equation approach, the dependence of entropy production has been studied in the works [5– 9]. For continuous systems, via the Fokker-Planck equation, several thermodynamics relations have been derived [11, 12, 17, 27, 28]. The method of calculating entropy production and extraction rates at the ensemble level by first analyzing thermodynamic relations at the trajectory level was introduced in the work [10]. Alternatively, many thermodynamic relations that derived via stochastic thermodynamics were reconfirmed under time reversal operation [29, 30]. At a single trajectory level, various thermodynamics relations have been derived via the fluctuation theorem [31–33]. The recent work [34] also indicates that for non-Markovian systems, the entropy production rate can be negative. All of these studies have explored entropy production based on full dynamical equations. However, inferring the entropy production for short experimental data is also vital but challenging since such studies lack a complete picture of the nature of interactions. In this regard, the recent fundamental relation called the thermodynamic uncertainty relation (TUR) sheds light on obtaining the expression for the entropy production from a given time series data [35– 38]. Recently, TUR has been generalized to study the entropy production for periodically driven systems [39].

Most recently, considering a single thermal ratchet, the general expressions for free energy, entropy production, as well as entropy extraction rates, are derived for a system that is genuinely driven out of equilibrium by time-independent force as well as by spatially varying thermal background [40]. However, protein-based molecular motors such as kinesin, myosin, and dynein walk in complex networks. Nanorobots are artificial Brownian motors and they are designed to achieve unidirectional motion on two-dimensional surfaces. Most physically important problems such as impurities (donors) diffuse along the semiconductor layer and they require 2D and 3D analysis.

In this work, extending our previous study [40], we further study the thermodynamic features of a single Brownian particle that walks along *M* Brownian ratchets arranged in a complex network. Each ratchet potential is either coupled with the hot and cold reservoirs or a heat bath where its temperature decreases linearly along the reaction coordinate. The analytical result depicts that the rate for the thermodynamic quantities such as the velocity V, entropy production *ė*_*p*_(*t*) as well as entropy extraction 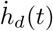 are independent of the network size at steady state as reconfirmed by complex generating functions. On the contrary, the thermodynamic relations such as the entropy S, entropy production *e*_*p*_(*t*), and entropy extraction *h*_*d*_(*t*) of the system step up with the network size*M* even at a steady state. We then further, study the system by including heat transfer via kinetic energy. Since the heat exchange via kinetic energy does not affect the energy dissipation rate, the heat dumped to the cold reservoirs contributes only to the internal entropy production. This implies that such a system exhibits a higher degree of irreversibility as long as a distinct temperature is retained between the hot and cold heat baths. The thermodynamic properties of systems that operate between the hot and cold baths are also compared and contrasted with a system that operates in a heat bath where its temperature varies linearly along with the reaction coordinate. Regardless of the network size *M*, the entropy, entropy production, and extraction rates are considerably larger for the linearly varying temperature case than a system that operates between the hot and cold baths revealing such systems are inherently irreversible.

We want to emphasize that a higher entropy is observed for a linearly decreasing temperature case than a Brownian particle that operates between the hot and cold baths. Consequently, systems that operate in a linearly decreasing temperature heat bath, have very low efficiency but a higher velocity. Thus such a thermal arrangement is beneficial in designing a device that channels a Brownian particle fast along the reaction coordinate. A Brownian particle that operates between two heat baths has a higher efficiency but a lower velocity in comparison with a linearly decreasing temperature case. Thus such a model system is advantageous if the sole purpose is to design an efficient Brownian motor. For all thermal arrangements, the particle attains a faster velocity whenever the network size *M* increases. This suggests that by properly arranging the model ingredient prior to the motor operation, a specific task can be accomplished. Thus the exactly solvable model presented in this work helps in designing artificial Brownian motors such as nanorobots.

The rest of the paper is organized as follows: in Section II, we present the model. The role of time is explored for a Brownian particle moving on M Brownian ratchets arranged in a complex network where each ratchet potential is coupled with the hot and cold baths. The model system is further explored by including heat transfer via kinetic energy. An alternative derivation for steady-state thermodynamic quantities via complex generating functions is also presented. In section III, we study the thermodynamic relations for Brownian particle that operates on M Brownian ratchets are arranged in a complex network. Each ratchet potential is coupled with a heat bath where its temperature decreases linearly along the reaction coordinate. Section IV deals with the summary and conclusion.

## II. BROWNIAN PARTICLE OPERATING IN A NETWORKS THAT COUPLED WITH THE HOT AND COLD RESERVOIRS

### A. The model

The Brownian particle undergoes a biased random walk along a discrete ratchet potential with load that coupled with the temperature

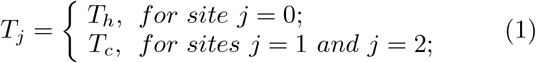

as shown in Figs. 1a and 1b. The potential barrier height *E >* 0, *f* denotes the load and *j* is an integer. *T*_*h*_ and *T*_*c*_ represent the temperature for the hot and cold reservoirs, respectively. The lattice has spacing *d* and in one cycle, the particle walks a net displacement of three lattice sites.as shown in Fig. 1. The potential *E >* 0, *f* denotes the load and *i* is an integer that runs from −∞ to ∞. *T*_*h*_ and *T*_*c*_ designate the temperature for the hot and cold reservoirs, respectively. Moreover, *d* denotes the lattice spacing *d* and in one cycle, the particle walks a net displacement of three lattice sites as shown in Fig. 1. The jump probability from site *i* to *i* + 1 is given by 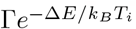 where Δ*E* = *U*_*i*+1_ − *U*_*i*_ and Γ is the probability attempting a jump per unit time. *k*_*B*_ designates the Boltzmann constant and hereafter *k*_*B*_, Γ and *d* are considered to be a unity. Obeying the metropolis algorithm when Δ*E ≤* 0, the jump definitely takes place while Δ*E >* 0 the jump takes place with probability exp(− Δ*E /T*_*i*_) [18].

**FIG. 1:**
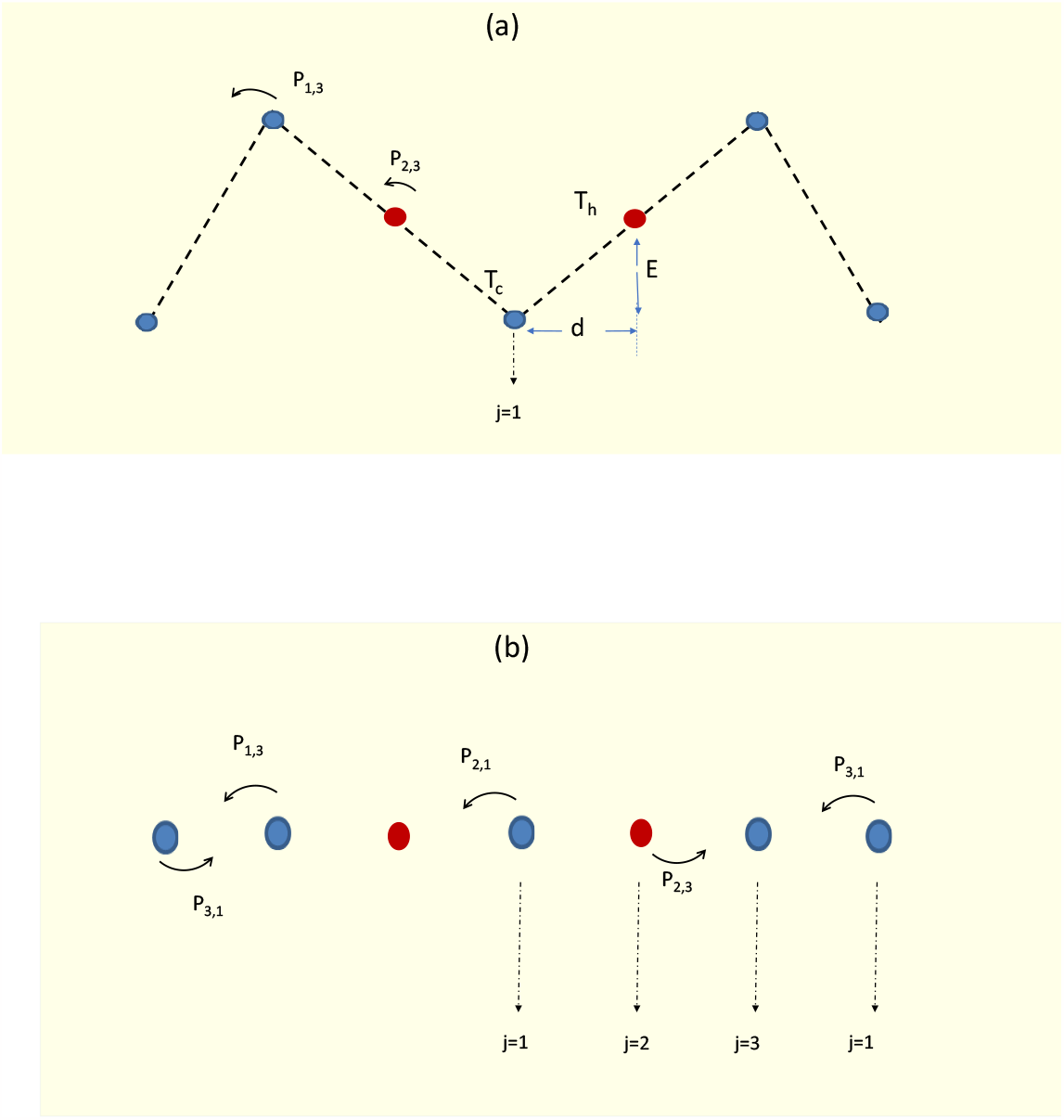
(Color online) (a) Schematic diagram (Side view) for a Brownian particle walking in two discrete ratchet potentials with the load. Sites with red circles are coupled to the hot reservoir (*T*_*h*_) while sites with blue circles are coupled to the cold reservoir (*T*_*c*_). Site 1 is labeled explicitly and *d* is the lattice spacing. (b) A Brownian particle (top view) walking in two discrete ratchet potentials with a load. Sites with red circles are coupled to the hot reservoir (*T*_*h*_) while sites with blue circles are coupled to the cold reservoir (*T*_*c*_).

It is also of practical interest to investigate the thermodynamic feature of *M* ratchet potentials arranged in a lattice (each potential has a similar arrangement as shown in Figs. 2a and 2b). This is because most practical problems such as neural system, cardiac system as well as impurities (donors) diffusion along the semiconductor layer require 2D and 3D analysis. As shown in Fig. 2a, each ratchet potential shares a common hot bath and a cold bath at the other end. When a periodic boundary condition is imposed, such arrangements can be mapped into the network depicted in Fig. 2b. For instance, the geometry of the networks shown in Fig. 2b and Fig. 2a are essentially the same (when *M* = 4) if one binds the nodes A, B, C, D of Fig. 2a together (when a periodic boundary condition is imposed).

**FIG. 2:**
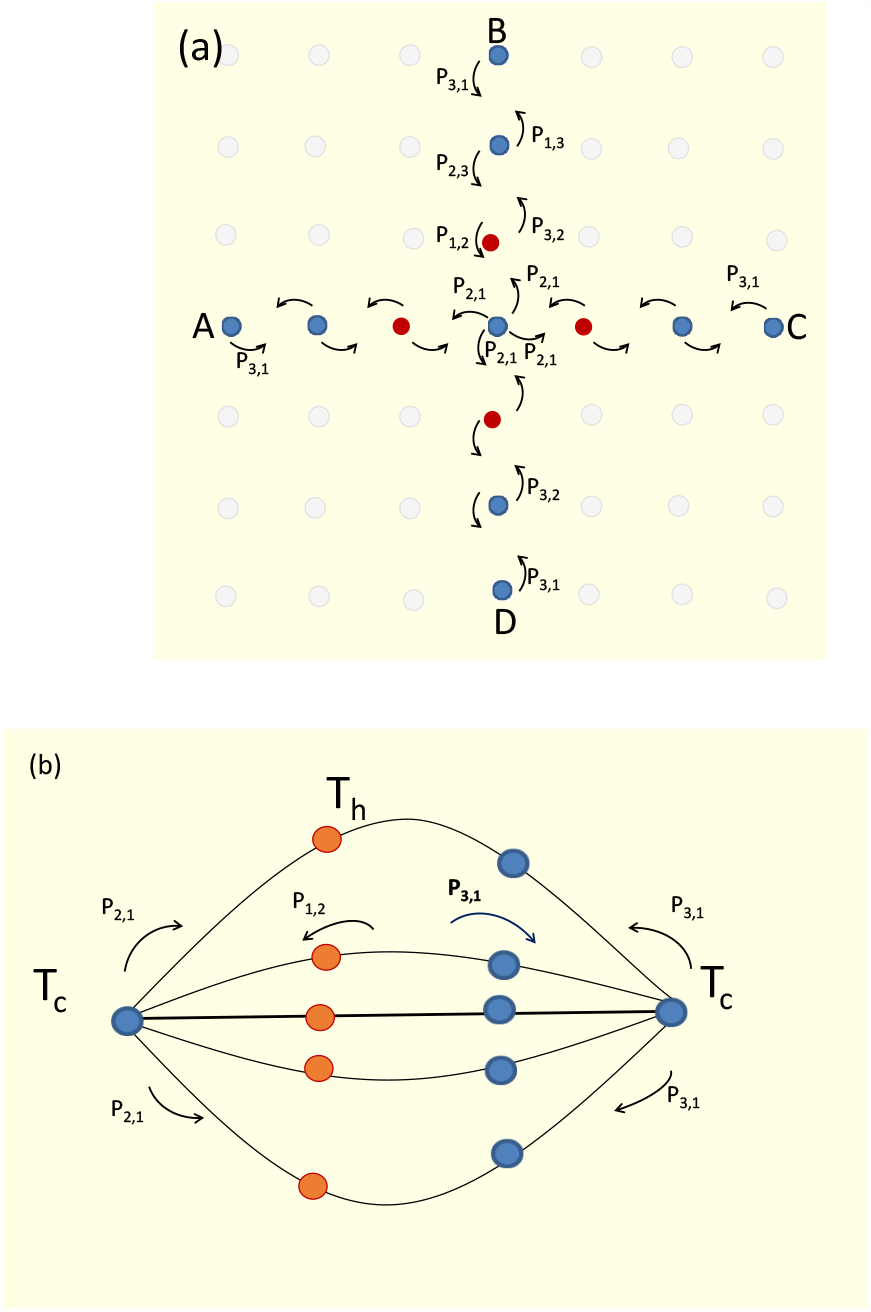
(Color online) (a) Top view of four thermal ratchets arranged in a lattice. Each ratchet potential has four lattice sites. Sites with red circles are coupled to the hot reservoir (*T*_*h*_) while sites with blue circles are coupled to the cold reservoir (*T*_*c*_). The hot bath for the four Brownian ratchets are located at the center. (b) *M* ratchet potentials arranged in a loop where each ratchet potentials share a common cold bath and a cold bath at the other end. Each ratchet potential has four lattice sites as shown in Fig. 2. Sites with red circles are coupled to the hot reservoir (*T*_*h*_) while sites with blue circles are coupled to the cold reservoir (*T*_*c*_). The hot bath for the *M* Brownian ratchets are located on the left side. The geometry of networks shown in Fig. 2a and Fig. 2b are essentially the same (when *M* = 4) if one binds the nodes A, B, C, D of Fig. 6b together (when a periodic boundary condition is imposed).

For a Brownian particle that moves in a discrete lattice, its dynamics is governed by the master equation

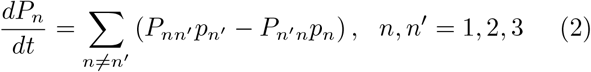

where *P*_*n′n*_ is the transition probability rate at which the system, originally in state *n*, makes a transition to state *n*^*′*^. *P*_*n′n*_ is given by the Metropolis rule [22]. Equation (2) can be rewritten as

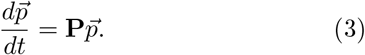

The expression for **P** can be evaluated for any *M*. For instance, for a single ratchet potential *M* = 1, **P** is a 3 by 3 matrix which is given by

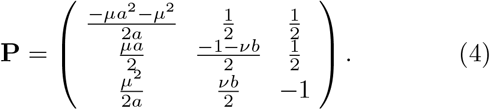

For two ratchet potentials (*M* = 2), **P** has a form

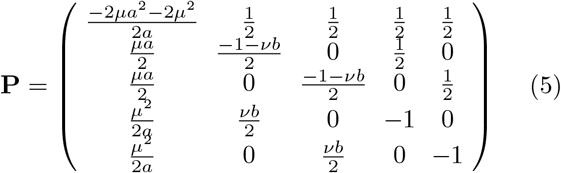

while for *M* = 3 and *M* = 4 case, one finds

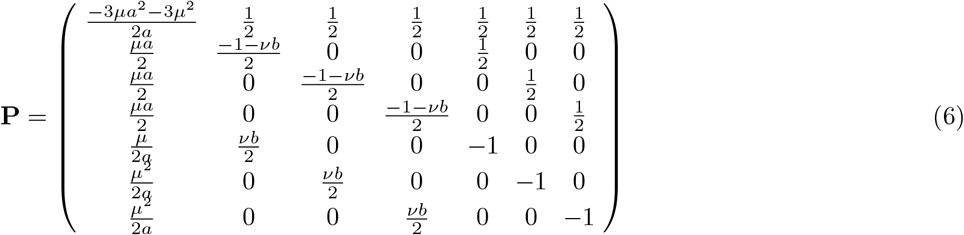

and

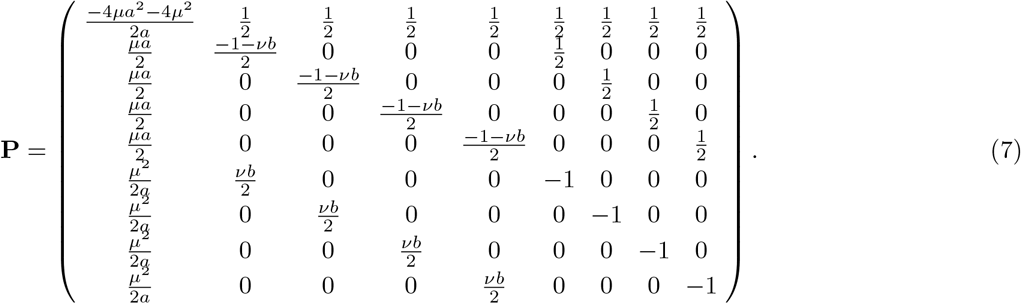

Here 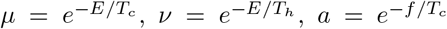 and 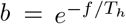. In Appendix A, the expressions for the probability are presented for the cases *M* = 1 and *M* = 2. The escaping rates

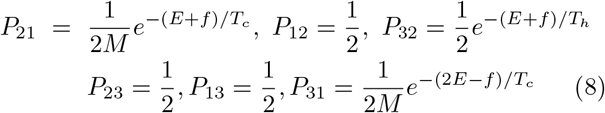

depend on *M*.

### B. Derivation of free energy

For a Brownian particle that moves along one or two dimensional discrete ratchet potential, the Boltzmann-Gibbs entropy relation

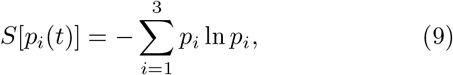

remains valid even when the system is far from equilibrium. The entropy extraction rate 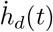 can be rewritten in terms of local probability density and transition probability rate as

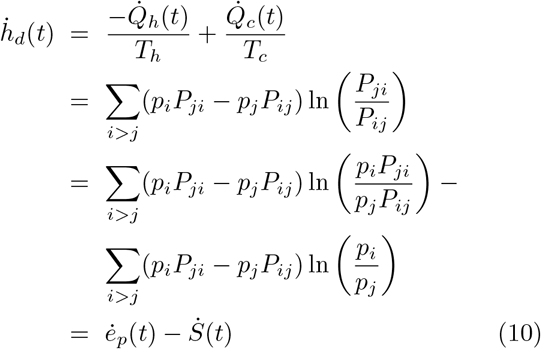

where

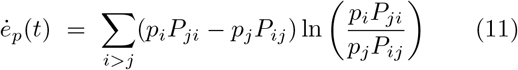

and

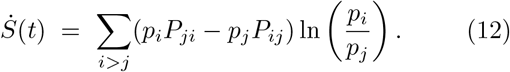

Here *ė*_*p*_(*t*) and 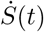 designate the entropy production rate and the rate of total entropy, respectively while 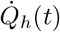 and 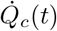 denote the heat per unit time taken from the hot reservoir and the heat per unit time given to cold reservoir, respectively.

After calculating the expressions for 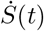, *ė*_*p*_(*t*) and 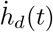 as a function of *t*, the analytic expressions for the change in entropy production, heat dissipation and total entropy can be found analytically via 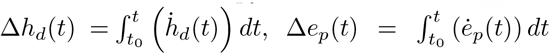 and 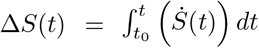 where Δ*S*(*t*) = Δ*e*_*p*_(*t*) *−* Δ*h*_*d*_(*t*).Once again, in the absence of load *f*, in the limit *T*_*h*_→ *T*_*c*_ and when *E* = 0, for both cases, one finds Δ*h*_*d*_(*t*) = 0,

In terms of local probability density and transition probability, the heat dissipation rate can be given as

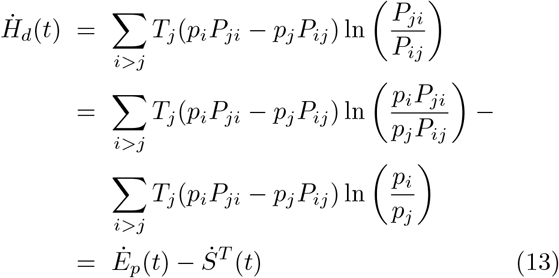

where

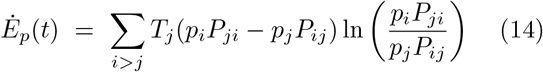

and

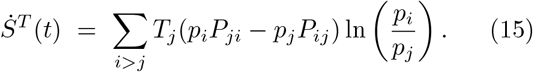

Our next objective is to write the expression for the free energy in terms of *Ė*_*p*_(*t*) and 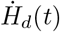 where *Ė*_*p*_(*t*) and 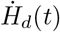 are the terms that are associated with *ė*_*p*_(*t*) and 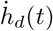 except the temperature *T*_*j*_.

The second law of thermodynamics can be written as Δ*S*^*T*^ (*t*) = Δ*E*_*p*_(*t*) − Δ*H*_*d*_(*t*) where Δ*S*^*T*^ (*t*), Δ*E*_*p*_(*t*) and Δ*H*_*d*_(*t*) are very lengthy expressions which can be evaluated via 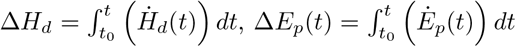 and 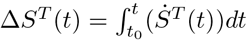. On the other hand, the total internal energy *U* (*t*) is the sum of the internal energies

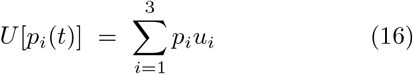

while the change in the internal energy is given by Δ*U* (*t*) = *U* [*p*_*i*_(*t*)] − *U* [*p*_*i*_(0)]. We also verify the first law of thermodynamics

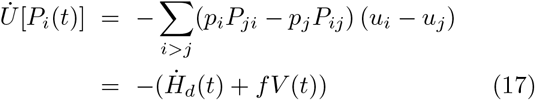

where the net velocity *V* (*t*) at any time *t* is the difference between the forward 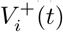 and backward 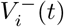 velocities at each site *i*. Next let us find the expression for the free energy dissipation rate 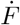. For the isothermal case, the free energy is given by *F* = *U* − *TS* and next we adapt this relationship to nonisothermal case to write

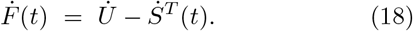

Substituting Eqs. (10) and (17) in Eq. (18) leads to

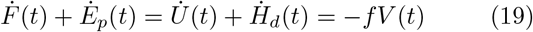

which is the second law of thermodynamics. Note that in the absence of load, 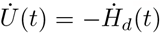 and consequently 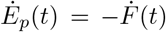. The change in the free energy can be written as

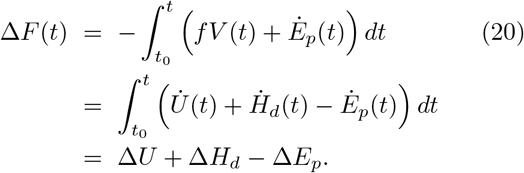

### C. Entropy, entropy production and extraction rates

The net velocity *V* (*t*) at any time *t* is the difference between the forward 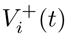 and backward 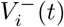 velocities at each site *i*. For instance, for *M* = 2 cases, the velocity can be calculated as

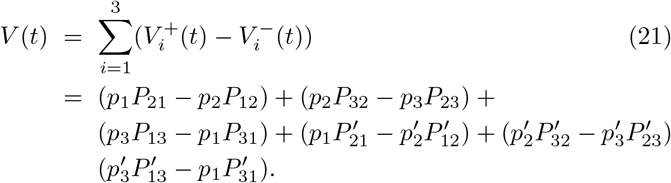

Because closed-form expressions for 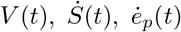 and 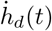 as a function of *t* is obtained, one can explore the dependence of other physical quantities as a function of time. To increase the readability of this paper, only the thermodynamic expressions up to *M* = 4 will be shown in this paper.

Hereafter, whenever we plot the figures, we use the following dimensionless load *λ* = *f/T*_*c*_, barrier height *ϵ* = *E/T*_*c*_ and dimensionless temprature *τ* = *T*_*h*_*/T*_*c*_.

#### The velocity

The dependence of the velocity on model parameters is explored as a function of time. The time *t* dictates the magnitude and direction of the velocity. For small *t*, the net particle flow is in the reverse direction (negative). As time increases, the direction of *V* is reversed. In Fig. 3a, we plot the velocity *V* as a function of *ϵ*. The figure depicts that the particle manifests a peak velocity at a particular barrier height *ϵ*^*max*^. At this particular height, the engine operates with maximum power. Phase space at which the velocity *V* = 0 is plotted in Fig. 3b as a function of *λ* and *τ*. In the figure, the potential height is fixed as *ϵ* = 8.0, *ϵ* = 6.0, *ϵ* = 4.0, and *ϵ* = 2.0 from top to bottom.

**FIG. 3:**
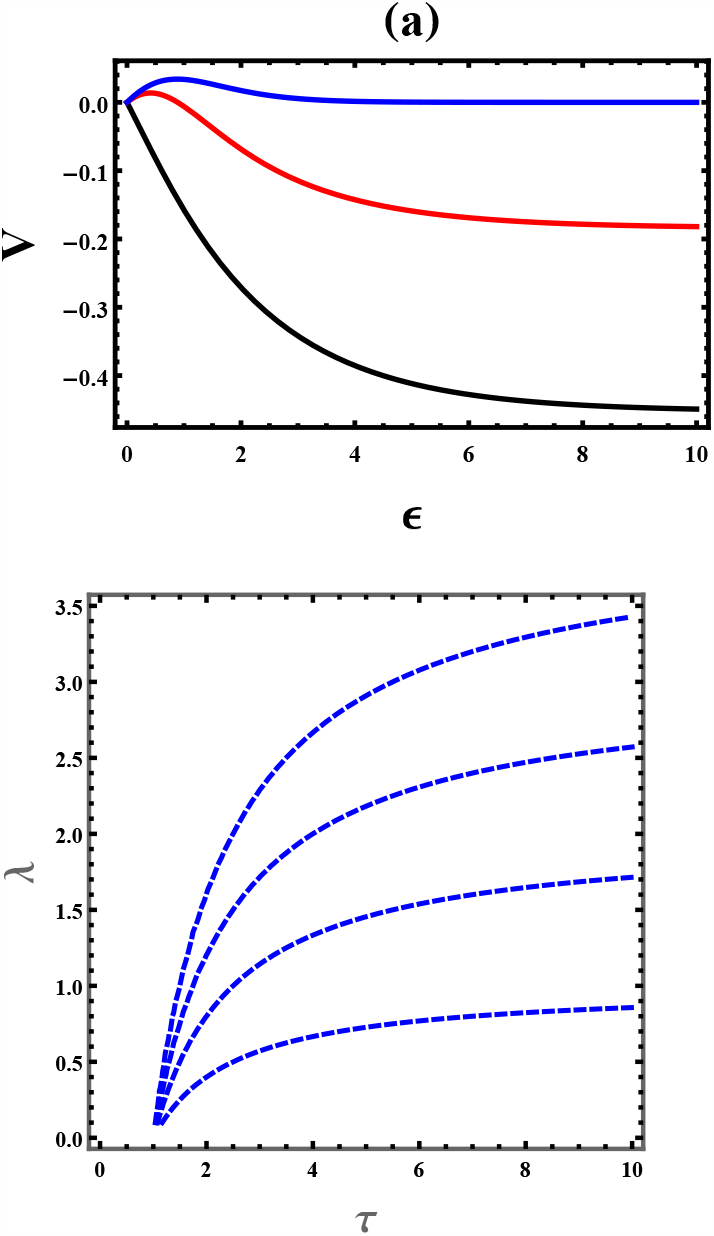
(Color online) (a) Particle velocity *V* as a function of *ϵ* is plotted via Eq. (21) for fixed *M* = 4 and *τ* = 2.0. The load is fixed as *λ* = 0, *λ* = 0.5, and *λ* = 1.0 from the top to bottom. (b) Phase space at which the velocity *V* = 0 is plotted as a function of *λ* and *τ*. The potential height is fixed as *ϵ* = 8.0, *ϵ* = 6.0, *ϵ* = 4.0 and *ϵ* = 2.0 from top to bottom.

The velocity increases as the number of ratchet potentials (*M*) steps up. At a steady state, the velocity approaches the same value regardless of the number of ratchet potentials arranged in the lattice. At steady state, for all *M*, the velocity approaches

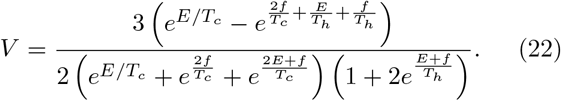

At stall force

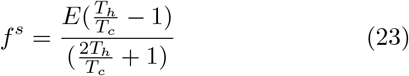

the velocity approaches zero. At this point, we want to emphasize that the model system not only serves as an important tool to calculate the velocity for thermally driven motors but also for isothermal motors (protein-based molecular motors) such as Myosins, Kinesin, and Dynein. When the motor operates in isothermal baths, the engines undergo unidirectional motion by utilizing chemical energy. For the isothermal case *T*_*h*_ = *T*_*c*_, the load only acts as the symmetry-breaking field. Thus in the presence of load, the motor attains a unidirectional velocity depending on the magnitude of the load.

#### The total entropy

The fundamental entropy relation

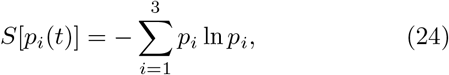

for the system which is far from equilibrium can be derived by substituting 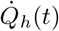 and 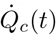 into Eq. (10). Here care must be taken that the change in the entropy Δ*S* = Δ*e*_*p*_ − Δ*h*_*d*_ reflects the system properties while the entropy production Δ*e*_*p*_ signifies the spontaneous change that the system undergoes. The dependence of entropy on the model parameter can be explored via Eq. (9). As shown in Fig. 4, the entropy of the system exhibits an intriguing parameter dependence. For *t* ≠ 0, *S >* 0 which indicates that in the presence of symmetry-breaking fields such as nonuniform temperature or external force, the system is driven out of equilibrium. *S*(*t*) is also considerably larger when four ratchet potentials are arranged (*M* = 4) in the network. The entropy decreases as the number of ratchet potentials (*M*) decreases which is reasonable since the entropy of the system depends on the number of acceptable states (lattices).

**FIG. 4:**
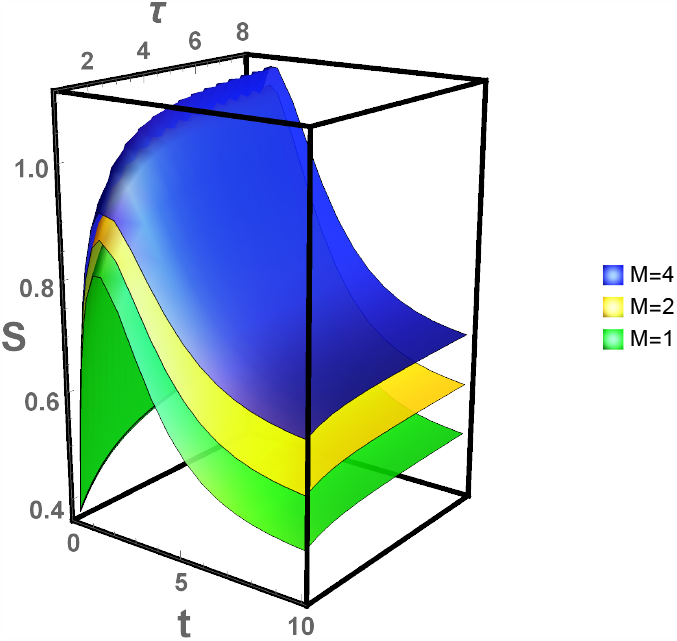
(Color online) The entropy *S*(*t*) as a function of *t* and *τ* evaluated analytically via Eq. (9) for a given *ϵ* = 2.0 and *f* = 0.0. In the figure, *M* is fixed as *M* = 4, *M* = 2, and *M* = 1 from top to bottom, respectively.

In the absence of load and at a steady state, the entropy approaches

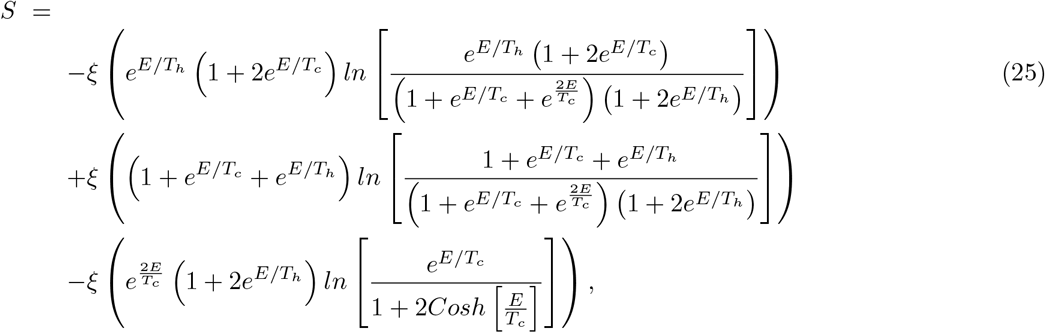

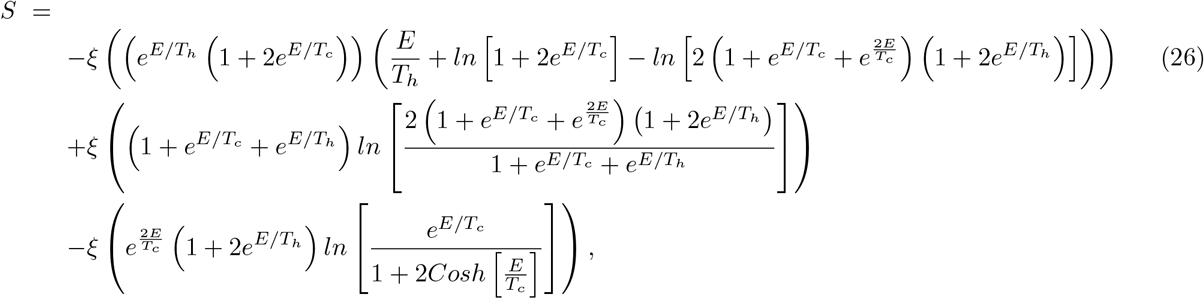

and

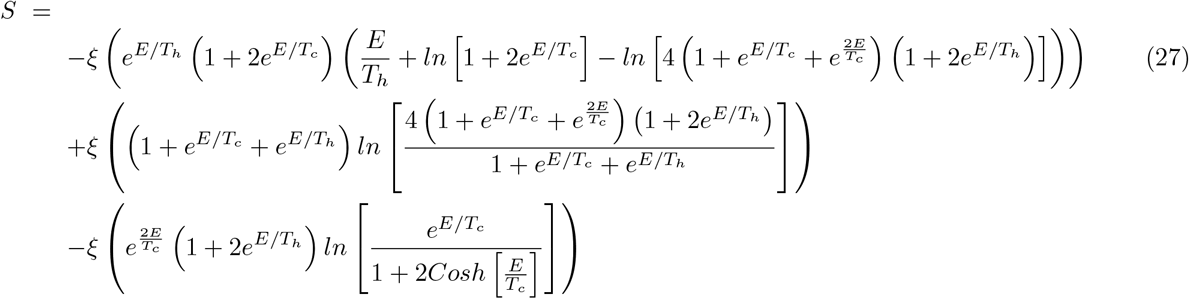

for *M* = 1, *M* = 2 and *M* = 4, respectively. Here 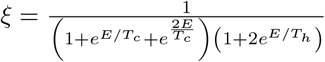.

Taylor expanding Eqs. (25), (26), and (27) near *E* = 0 leads to

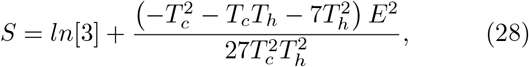

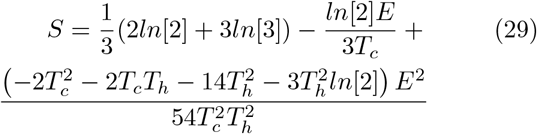

and

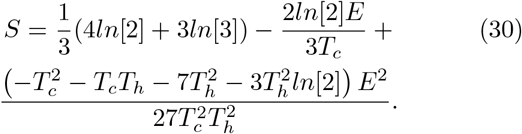

At equilibrium, when *E*→ 0, Eqs. (28), (29), and (30) converge to *S* = *ln*[3], *S* ≈ *ln* [5] and *S* ≈ *ln*[9] which reasonable since *S* relies on the lattice size. The system can be driven out of equilibrium as long as it operates at a finite time even in the absence of potential and load. This can be appreciated by analyzing the general time-dependent entropy *S* in the limit *f* → 0, *T*_*h*_→ *T*_*c*_, and *E*→ 0. Near *t* = 0, we get *S* = (1 + *ln*[2] − *ln*[*t*])*t* for all *M* and in the limit *t* → ∞, we recover *S* = *ln*[3], *S ≈ ln*[5] and *S* ≈ *ln*[9] for *M* = 1, *M* = 2 and *M* = 4, respectively.

#### Entropy production and extraction rates

In this section, the dependence for the rate of entropy production *ė*_*p*_(*t*), rate of entropy 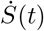, and rate of entropy extraction 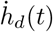 on the system parameters will be explored. Via Eqs. (13), (14), and (15), the expression for 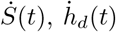 and *ė*_*p*_(*t*) can be evaluated. Fig. 5 shows the plot of 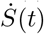 as a function of function of *t, τ* and *M*. As depicted by the figure, the rate of entropy 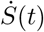 is higher for larger lattice size *M*. Fig. 6 shows the plot of *ė*_*p*_(*t*) as a function of *t, τ* and *M*. *ė*_*p*_(*t*) significantly large when *M* is large. The plot of 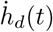 as a function of *t, τ*, and *M* is exhibited in Fig. 7. The figure shows that 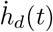 significantly becomes large as the network size steps up. The three figures also indicate that *ė*_*p*_(*t*) and 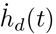 approach their steady state values 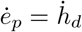 as time progresses. Our analysis also confirms that in the presence of symmetry-breaking fields, the entropy production rate *ė*_*p*_ steps down in time and at steady state, 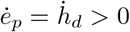. However, for an isothermal case and in the absence of load, 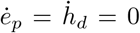 in a long time limit.

**FIG. 5:**
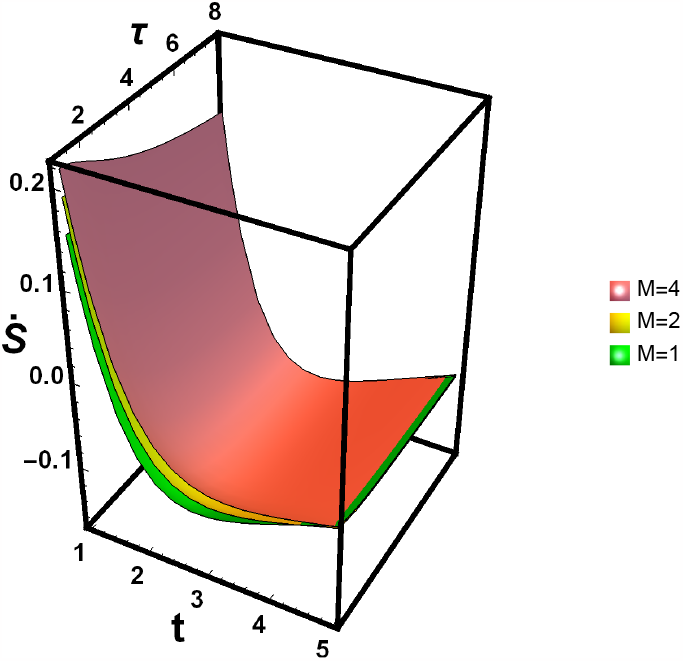
(Color online) 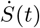 versus *t* is evaluated analytically for fixed *ϵ* = 2.0, *τ* = 2.0 and *λ* = 0.6. In the figure, *M* is fixed as *M* = 4, *M* = 2, and *M* = 1 from top to bottom respectively. As shown in the figure, 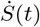decreases as *M* and *t* increase. At a steady state, all of these thermodynamic quantities approach the same value.

**FIG. 6:**
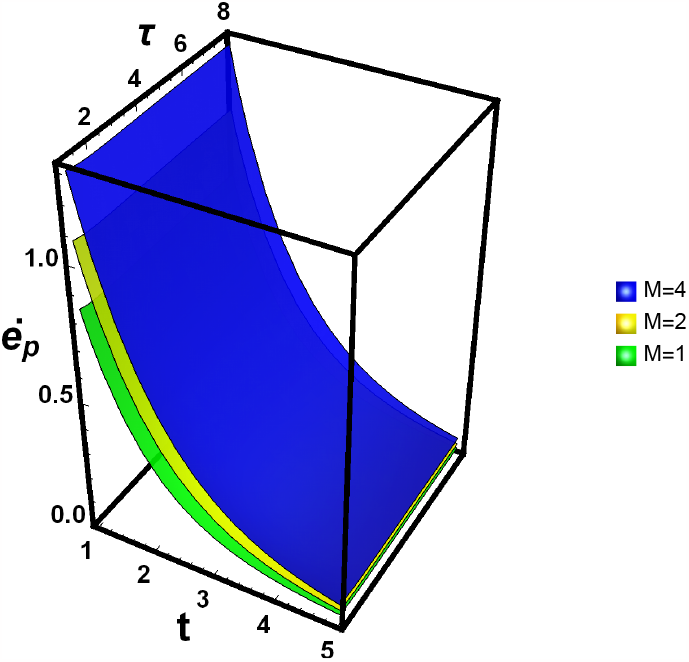
(Color online) The entropy production rate *ė*_*p*_(*t*) as a function of *t* for fixed *ϵ* = 2.0, *τ* = 2.0 and *λ* = 0.6. *M* is fixed as *M* = 4, *M* = 2 and *M* = 1 from top to bottom, respectively. As shown in the figure, *ė*_*p*_(*t*) decreases as *M* and *t* increase. At a steady state, all of these thermodynamic quantities approach the same value.

**FIG. 7:**
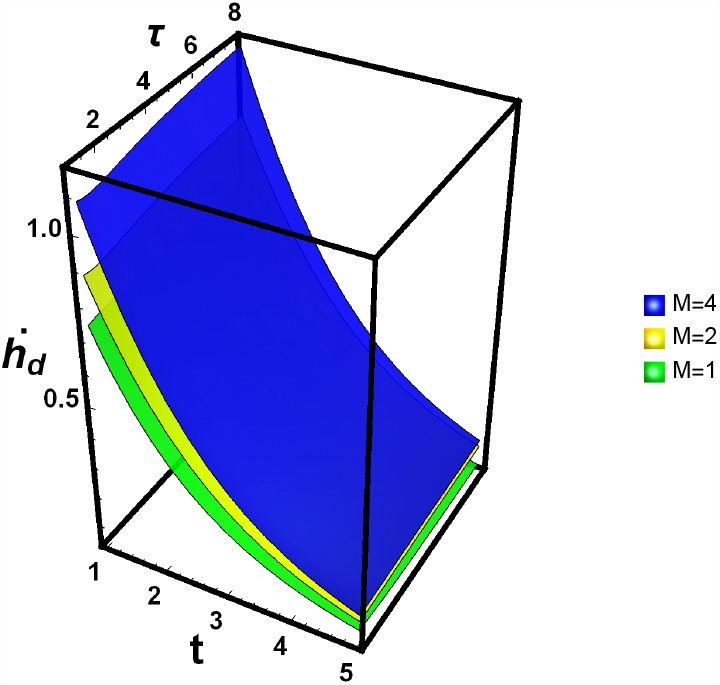
(Color online) The entropy extraction rate 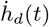 as a function of *t* evaluated analytically using Eq. (3) for a given value of *ϵ* = 2.0, *τ* = 2.0, *λ* = 0.6. In the figure, *M* is fixed as *M* = 4, *M* = 2, and *M* = 1 from top to bottom respectively. As shown in the figure, 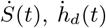 and *ė*_*p*_(*t*) decrease as *M* and *t* increase. At a steady state, all of these thermodynamic quantities approach the same value.

For any *M*, at steady state we find

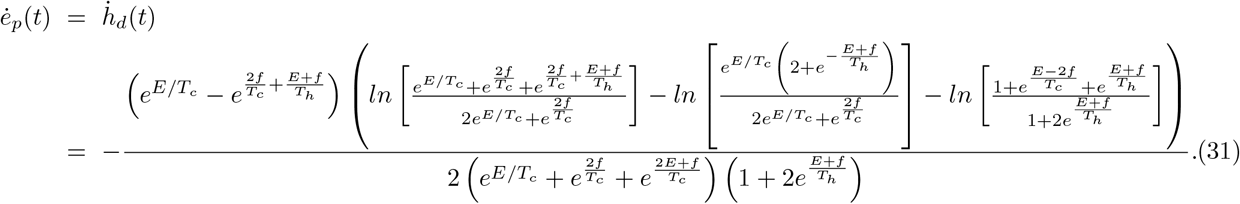

Analyzing the entropy production is not only vital in describing non-equilibrium systems but also signifies the degree of irreversibility of a given system. As indicated by Eq. (31), 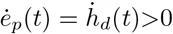 as long as *f* ≠ 0 and when a distinct temperature between the hot and cold bath is retained *T*_*h*_ ≠ *T*_*c*_. At stall force (in the limit *f* → *f*^*s*^), 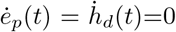. The phase space at which the entropy production rate becomes zero is depicted in Fig. 8.

**FIG. 8:**
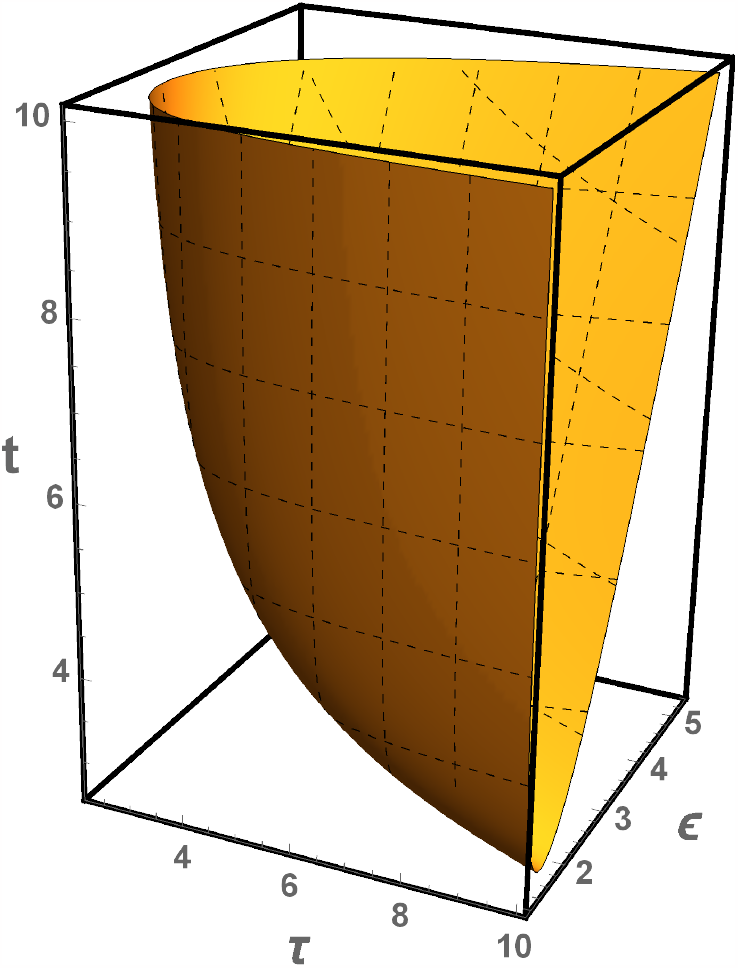
(Color online) (a) The Phase space at which the entropy production rate becomes zero for fixed *λ* = 0.6.

In the absence of load *f*, potential *E*, and in the limit *T*_*h*_ → *T*_*c*_, the system relaxes to its equilibrium state in the long time limit. On the contrary, in the short time limit, the system still operates irreversibly. This can be appricated by analyzing 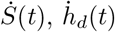 and *ė*_*p*_(*t*) in the absence of load *f*, potential *E*, and in the limit *T*_*h*_ → *T*_*c*_. For *M* = 1 case, one finds 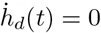 and

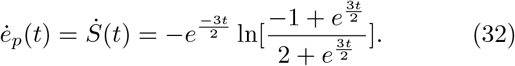

On the other hand, for *M* = 2 case, we get

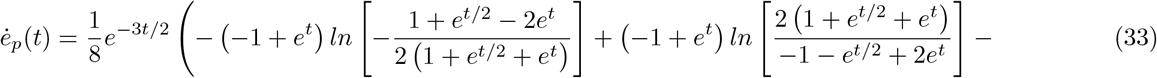

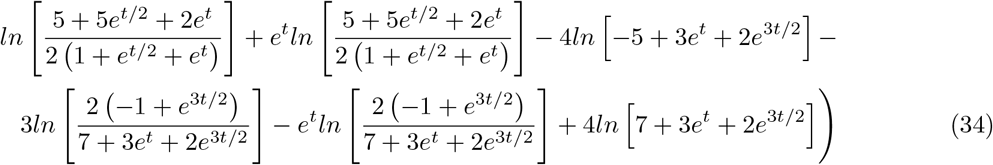

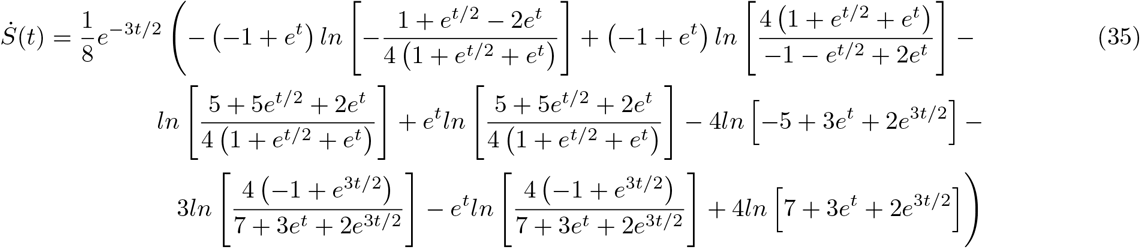

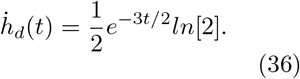

Once again in a quasistatic limit, regardless of any parameter choice, we find 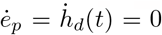 As pointed out by Ge *et. al*. [44], the vanishing of velocity may not indicate the system is at thermodynamic equilibrium

The the analytic expressions for the change in entropy production, heat dissipation, and total entropy can be found analytically via 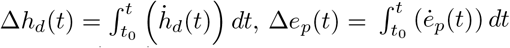and 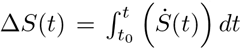 where Δ*S*(*t*) = Δ*e*_*p*_(*t*) − Δ*h*_*d*_(*t*) and the indexes *i* = 1*· · ·* 3 and *j* = 1*· · ·* 3. Unlike the entropy production and extraction rates, the thermodynamic relationship such as Δ*S*(*t*), Δ*e*_*p*_(*t*) and Δ*h*_*d*_(*t*) depend on network size even at steady state. The expressions for Δ*h*_*d*_(*t*), Δ*S*(*t*), and Δ*e*_*p*_(*t*) are lengthy and will not be presented in this work. As shown in Figs. 9a and 9b, for a system that operates between hot and cold reservoirs, Δ*e*_*p*_(*t*) and Δ*h*_*d*_(*t*) approach a non-equilibrium steady state in the long time limit. As the number of lattice size steps up, Δ*e*_*p*_(*t*) and Δ*h*_*d*_(*t*) increase showing for large *M* the system exhibits a higher level of irreversibility.

**FIG. 9:**
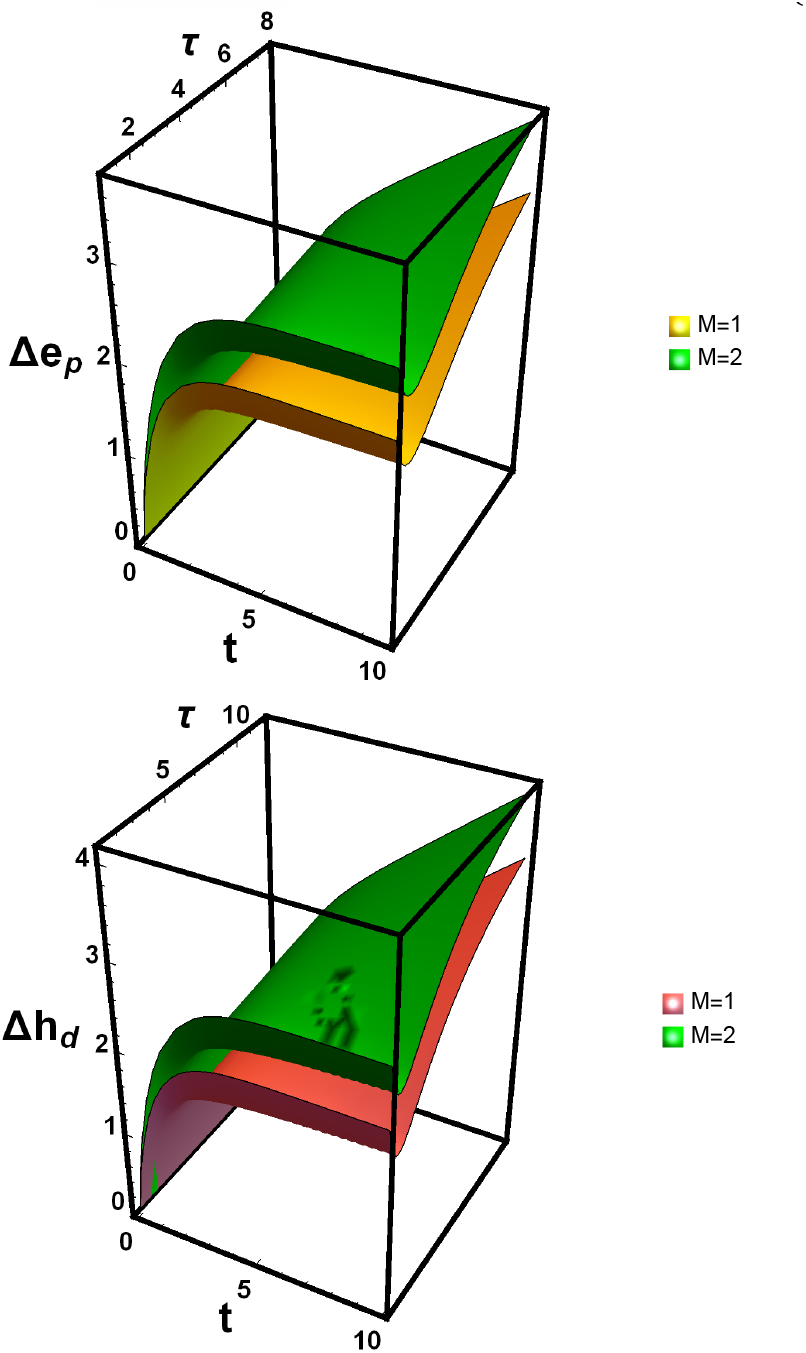
(Color online) (a) Δ*e*_*p*_(*t*) as a function of *t* and *τ* = 2.0 for fixed *ϵ* = 2.0 and *λ* = 0.0. (b) Δ*h*_*d*_(*t*) as a function of *t* and *τ* = 2.0 for fixed *ϵ* = 2.0 and *λ* = 0.0. In the figures, *M* is fixed as *M* = 4 and *M* = 1 from top to bottom respectively. As shown in the figures, Δ*e*_*p*_(*t*) and Δ*h*_*d*_(*t*) increase as *M* steps up and when *t* increases.

The second law of thermodynamics can be also written as Δ*S*^*T*^ (*t*) = Δ*E*_*p*_(*t*) − Δ*H*_*d*_(*t*) where Δ*S*^*T*^ (*t*), Δ*E*_*p*_(*t*) and Δ*H*_*d*_(*t*) are very lengthy expressions which can be evaluated via 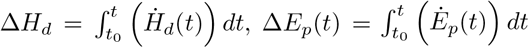 and 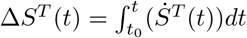.

As discussed before, once the expressions for 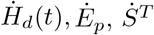 are analyzed, the corresponding entropy balance equation can be calculated as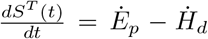. The expressions for these relations are very complicated. In Figs. 10a we plot Δ*E*_*p*_(*t*) as a function of *t* and *ϵ* for fixed *τ* = 2.0 and *λ* = 0. Fig. 10b depicts the plot of Δ*H*_*d*_(*t*) as a function of *t* and *τ* for fixed *ϵ* = 2.0 and *λ* = 0.6. In both figures, *M* is fixed as *M* = 2 and *M* = 1 from top to bottom respectively. As shown in the same figures, Δ*E*_*p*_(*t*) and Δ*H*_*d*_(*t*) increase as *M* and *t* increase.

**FIG. 10:**
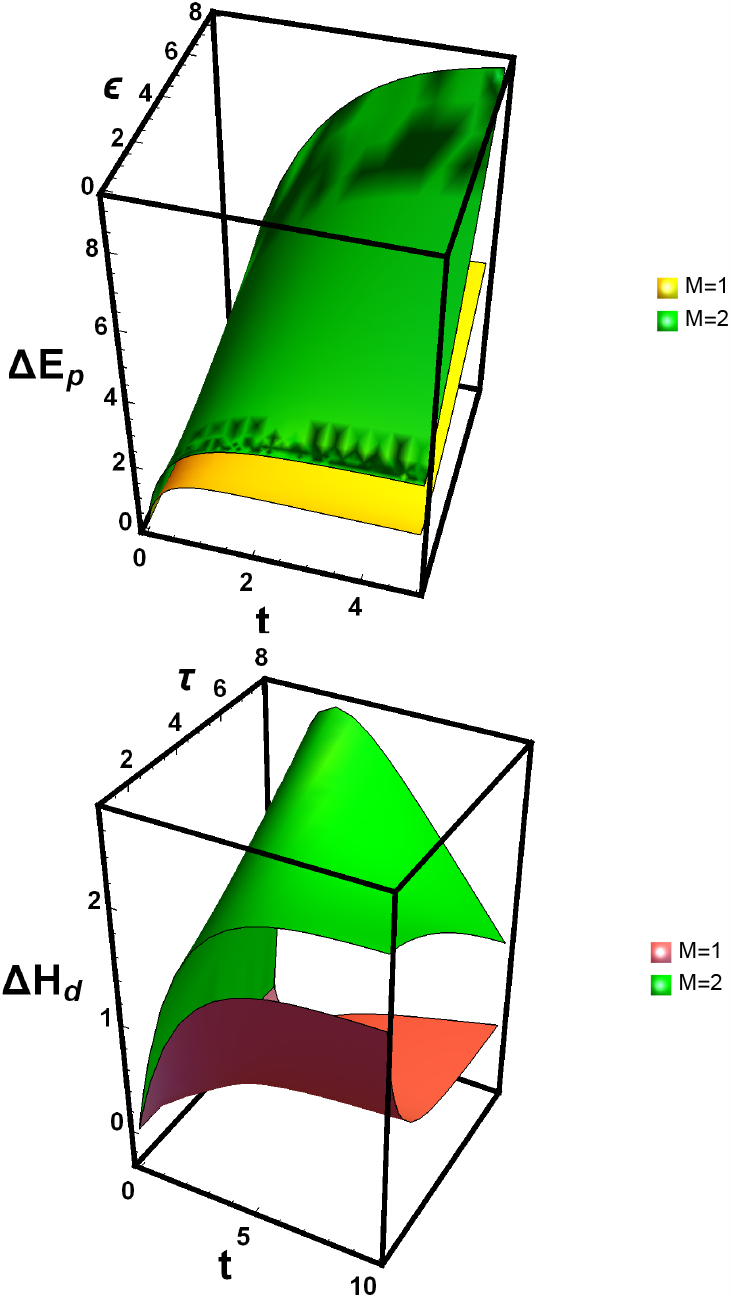
(Color online) (a) Δ*E*_*p*_(*t*) as a function of *t* and *ϵ* for fixed *ε* = 2.0 and *λ* = 0. (b) Δ*H*_*d*_(*t*) as a function of *t* and *τ* for fixed *ϵ* = 2.0 and *λ* = 0.6. In the figures, *M* is fixed as *M* = 2 and *M* = 1 from top to bottom respectively. As shown in the figures, Δ*E*_*p*_(*t*) and Δ*H*_*d*_(*t*) increase as *M* steps up and when *t* increases.

#### The free energy

Next, we explore the dependence of the free energy dissipation rate shown in Eqs. (12) and (13) on *t*. In general 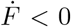 and approaches zero in the long time limit for both cases (see Fig. 11). As depicted in Fig. 11, when the system operates at finite time, the free energy dissipation rate increases as the number of ratchet potentials *M* steps up. At a steady state, the free energy rate approaches the same value for all *M*. This can be appreciated by evaluating the rate in the long limit. At a steady state, for all *M*, the free energy rate converges to

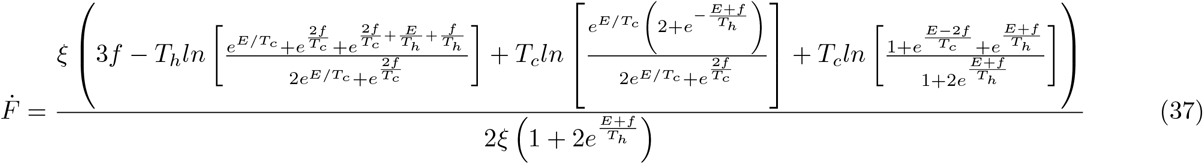

where 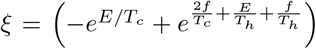. For the isothermal case, in the absence of potential barrier and load, Eq. (37) approaches zero since the detail balance condition is fulfilled. Moreover Δ*U* = Δ*H*_*d*_ = 0 and Δ*F* (*t*) = − Δ*E*_*p*_. At equilibrium (*t* →∞), we get Δ*F* = − *T*_*c*_ ln(3) for *M* = 1.

**FIG. 11:**
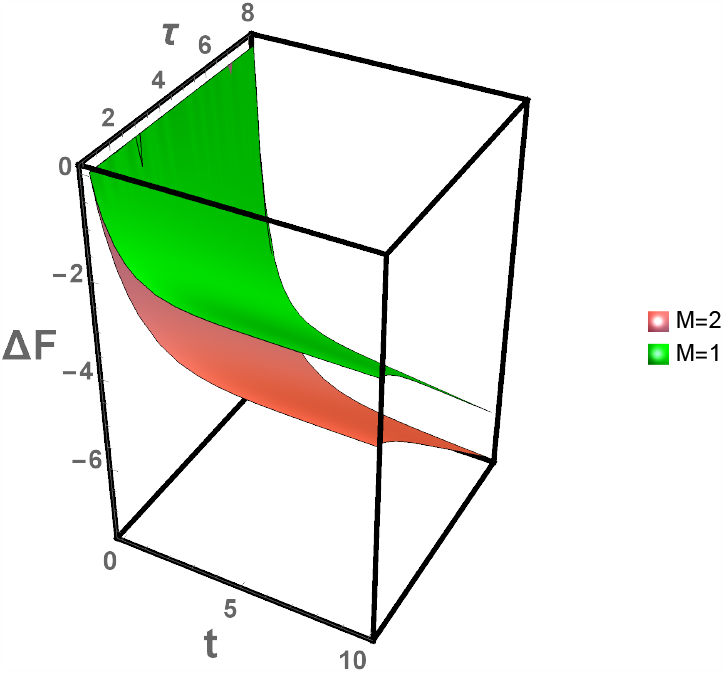
(Color online) (a) The change in free energy Δ*F* versus *t* and *τ* for *f* = 0.0 and *ϵ* = 2.0. In the figure, *M* = 1 and *M* = 2 from the top to bottom.

When the system operates at a finite time, the system operates irreversibly, in the absence of potential barrier and load. Taylor expanding the free energy rate near *t* = 0, one gets 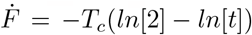 for *M* = 1 and 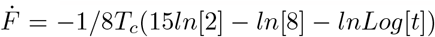 for *M* = 2 case. The change in free energy Δ*F* versus *t* and *τ* is depicted in Fig. 11 for fixed *f* = 0.0 and *ϵ* = 2.0. In the figure, *M* = 1 and *M* = 2 from the top to bottom. from the top to bottom.

### D. Heat transfer via kinetic energy

So far, the rate of heat loss due to particle recrossing at the boundary between the hot and cold reservoirs is not included. When the heat exchange via kinetic energy is included, the system becomes irreversible even at the quasistatic limit, and as a result *ė*_*p*_ *≠* 0 or *Ė*_*p*_ *≠*0.

During the biased random walk, the particle absorbs *k*_*B*_(*T*_*h*_− *T*_*c*_)*/*2 amount of heat from the hot bath. It then gives back *k*_*B*_(*T*_*h*_− *T*_*c*_)*/*2 amount of heat to the cold bath. As a result, there will be an irreversible heat flow from the hot to cold baths. The rate of heat transfer from the hot to the cold reservoirs 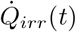 depends on how often the particle jumps from the cold to the hot heat baths. After some algebra, one gets

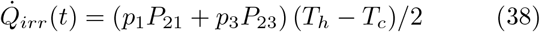

for *M* = 1 case and

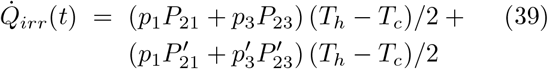

for *M* = 2 case. At steady state, for all *M* we find

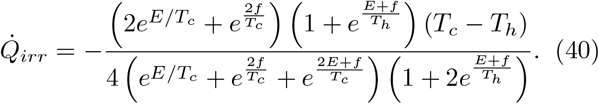

For the case where *f* = 0 and *E* = 0, one gets

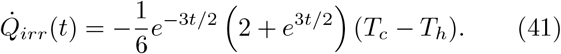

Integrating Eq. (41) with respect to time yields

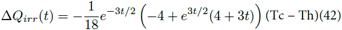

as long as *f* = 0 and *E* = 0.

One can note that the heat transfer through the kinetic energy does not affect 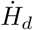 as the heat taken from the hot reservoirs goes to the cold reservoir.*This also implies that the whole heat dumped to the cold reservoirs contributes to the internal entropy production and hence for any parameter choice Ė*_*p*_(*t*) *>* 0 *as long as T*_*h*_ ≠ *T*_*c*_. *The heat loss due to particle recrossing only contributes to the internal entropy production and we infer the new entropy production rate to be*

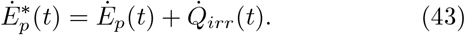

One can then rewrite the thermodynamic relations derived in the previous section in terms of 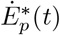. Here 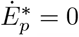 only when *T*_*h*_ = *T*_*c*_.

The rate of free energy depicted in Eq. (19), can be rewritten as

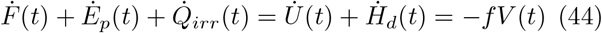

The change in the free energy is given by

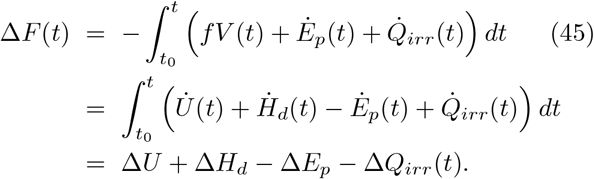

In the absence of load and potential energy, for all *M* at steady state, Eq. (44) converges to

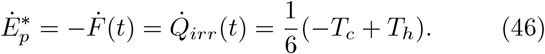

Eq. (46) confirms the second law of thermodynamics in essence that the entropy for a thermally driven heat engine is greater than zero. In other words, such systems operate irreversibly as long as a distinct temperature difference is retained between the hot and cold heat baths. For isothermal cases *T*_*h*_ → *T*_*c*_, 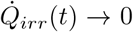 as expected. Eq. (46) also reconfirms Clausius’ statement of the second law [1] that *“Heat can never pass from a colder to a warmer body without some other change, connected therewith, occurring at the same time*.*”* In the next section, using complex generating functions, we show the rate for thermodynamic relations does not depend on the network size *M* at steady state.

### E. Alternative derivation of velocity via complex generating functions

In this section, we show why the rate for thermodynamic relations does not depend on the network size *M* at steady state, the mathematical approach discussed by Goldhirsch *et. al*. [45, 46] is adapted. Using this mathematical approach, we also calculated the velocity of the motor in our recent paper [47].

For clarity, first, we revise the mathematical technique developed. Consider a segment with sites 0, 1, 2....*N* as shown in Fig. 1b. Let *P*_2,1_ be the probability per unit time step for the walker to jump from 1 to 2 while *P*_1,2_ be the probability to jump from 2 to 1 with a condition *P*_1,2_ + *P*_2,1_ ≤ 1. The probability *P*_*w*_(*n*)^+^ designates the probability for the walker to start at *j* = 0 and reach *j* = *N* in *n* steps in the right direction. The mean first passage for the walker to reach *j* = *N* for the first time is given as [47]

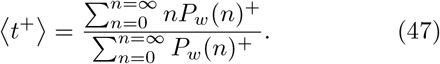

The corresponding generating function in terms of the *ϕ* probability for *P*_*w*_(*n*)^+^ is given by

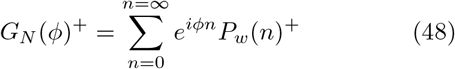

where in this case 0 *< ϕ <* 2*π*. In terms of generating function, the corresponding mean first passage time shown in Eq. (46) can be written as

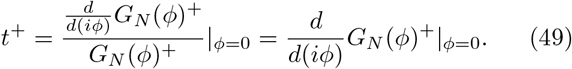

*P*_1_, *P*_2_, … are given by *P*_1_*e*^*iϕ*^, *P*_2_*e*^*iϕ*^, … in terms of *ϕ* probabilities. The particle has a probability (1− *P*_1_)^*n*^ to stay at *j* = 0 after *n* consecutive steps. The corresponding generating function is written as

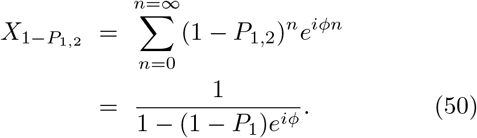

Similarly, in terms of generating function, the probability of staying at site *j* after *n* consecutive steps is given by

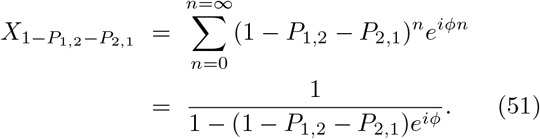

Let us *U* (*n*)^+^ to be the probability for the walker to leave *j* = 0 and reaches *j* = *N* in the right direction without returning to *j* = 0. The associated generating function can be expressed as

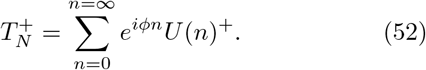

Similarly, let *V* (*n*)^+^ be the probability for the walker to leave *j* = 0 in the right direction and returns to *j* = 0 without reaching *j* = *N*. The corresponding generating function is given by

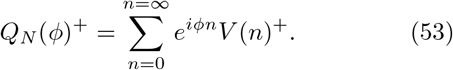

To reach the point *j* = *N*, the particle can stay at the site *j* = 0 for a number of steps with probability 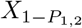 moves out, and returns to *j* = 0 without touching *j* = *N* with 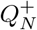 probability then stay at *j* = 0. These recursive steps are repeated many times as shown below

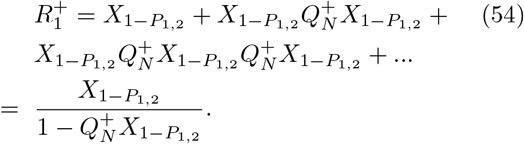

The particle finally will reach the site *j* = *N* without returning to *j* = 0 with the *ϕ* probability of 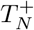 after several trials. Thus the generating function *G*_*N*_ (*ϕ*)^+^ for the first passage time takes a form

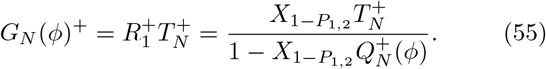

For the Brownian particle moving in complex networks, let us now calculate the generating function for the MFPT in the right direction (see Fig. 2a). To reach the other side of the potential well, the particle can stay at the site where the hot bath is located for a number of steps with probability 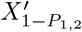, moves out and returns to the hot bath using one of the *M* possible paths with *ϕ* probability 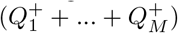. The particle then stays in the hot bath. The particle repeats these recursive steps many times as shown below

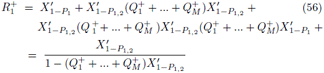

where 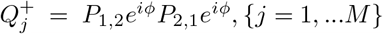 is defined as the *ϕ* probability for the walker to leave the hot bath and returns back to the hot bath without touching the other side of the potential well. After several trials, the particle finally will be able to reach the other side of the potential without returning to the hot bath using one of the possible *M* paths with *ϕ* probability 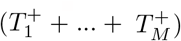. Here 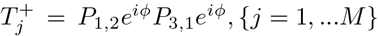 is the *ϕ* probability (for the *j*^*th*^ ratchet potential) for the walker to leave the hot bath and reach the other side of the potential well in the right direction without returning to the hot bath. The generating function for the MFPT towards the right direction is given by

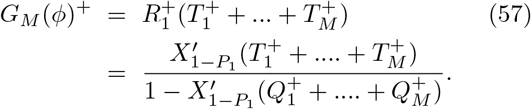

Following the above steps, the generating function for the MFPT to the left direction can be also given as

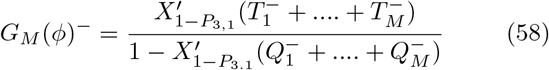

where 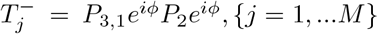 is the *ϕ* probability (for the *j*^*th*^ ratchet potential) for the walker to leave the cold bath (left potential well) and to reach the other side of the potential well in the left direction without returning to the cold bath while 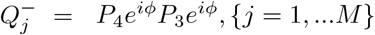 is defined as the *ϕ* probability for the walker to leave the cold bath (towards the left direction) and return to the cold bath without touching the other side of the potential well. For clarity let us write 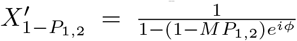 and 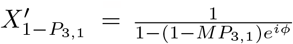. Care must be taken that now 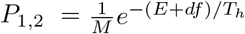, 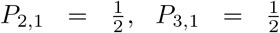 and 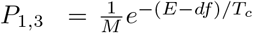. Since we have *M* identical potentials, the net average velocity (independent of lattice size *M*) is simplified to

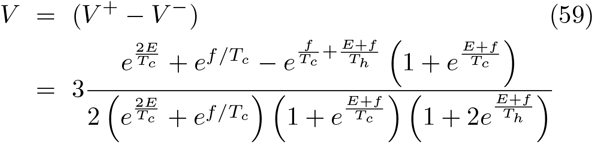

The approximated velocity in Eq. (59) is approximately the same as the velocity shown in Eq. (21). This can be appreciated by plotting *V* as a function of time *t* in Figs. 12a and 12b.

**FIG. 12:**
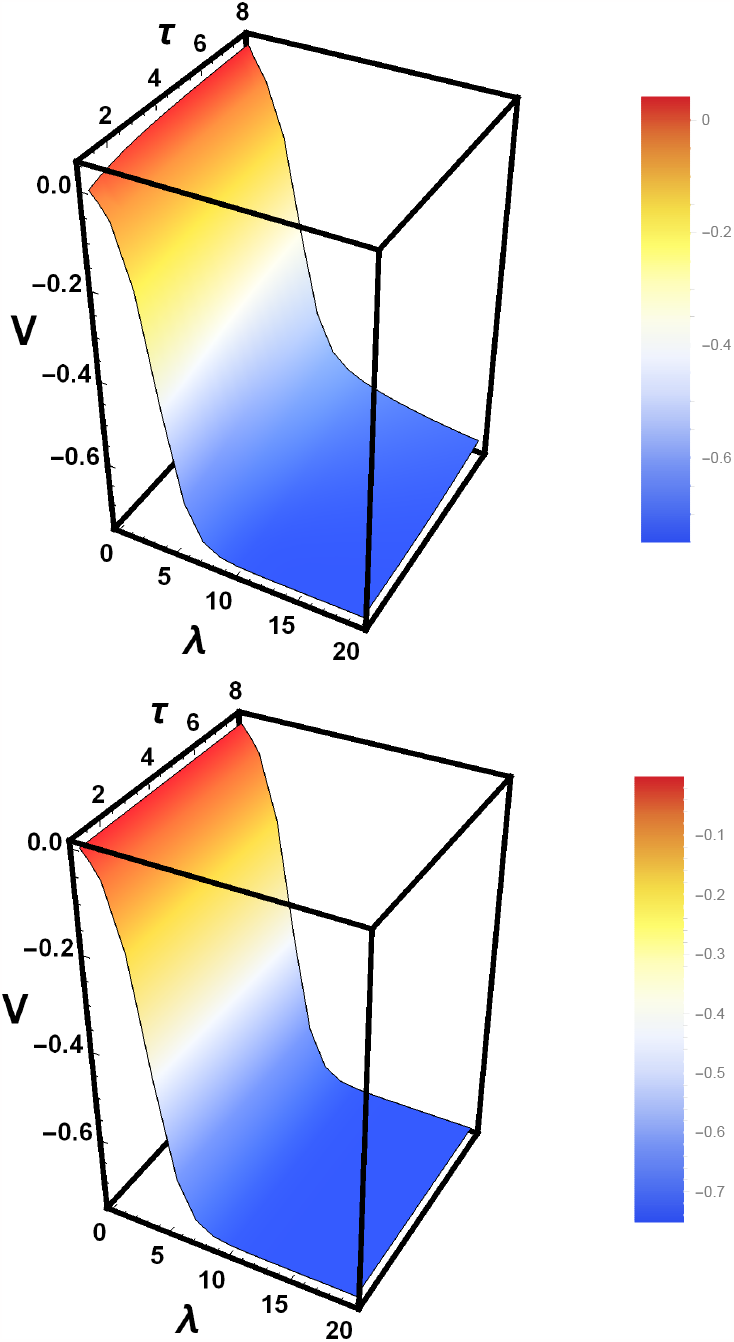
(Color online) (a) Particle velocity *V* as a function of *τ* and *λ* is plotted via Eq. (21) for fixed *ϵ* = 2.0. (a) Particle velocity *V* as a function of *τ* and *λ* is plotted via Eq. (59) for fixed *ϵ* = 2.0.

## III. BROWNIAN PARTICLE OPERATING IN A HEAT BATH WHERE ITS TEMPERATURE LINEARLY DECREASES ALONG WITH THE REACTION COORDINATE

Let us now consider *M* ratchet potentials arranged in a lattice (see Fig. 13) where each ratchet potential is coupled with a linearly decreasing temperature

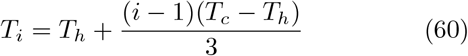

as shown in Fig. 14. Once again, the rate equation for the model is given by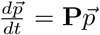 where the expression for **P** can be evaluated for any *M*. For instance, for a single ratchet potential *M* = 1, **P** is a 3 by 3 matrix which is given by

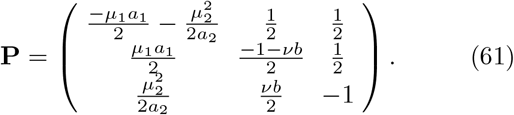

**FIG. 13:**
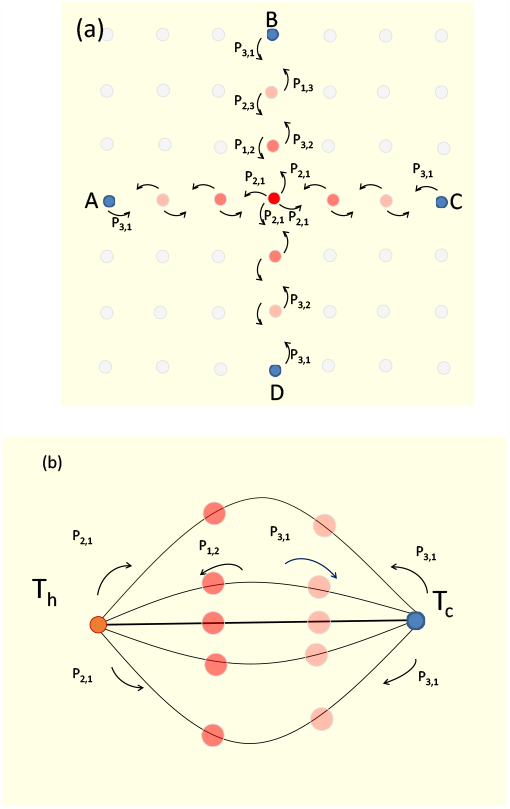
(Color online) (a) Four thermal ratchets are arranged in a lattice where in each ratchet potential the temperature is arranged to decrease linearly from the hotter bath (*T*_*h*_) to the colder bath (*T*_*c*_). The hotter bath for the four Brownian ratchets are located at the center. (b) A schematic diagram is plotted by imposing a periodic boundary condition (when one binds the nodes A, B, C, D of Fig. 13a together). Figs. 13a and 13b are the same (when *M* = 4). O

**FIG. 14:**
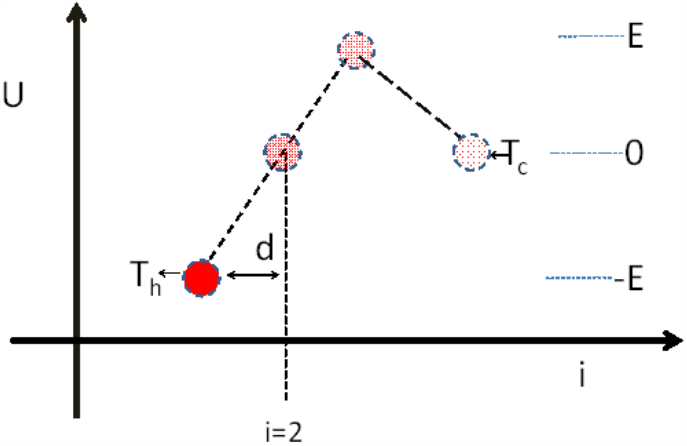
(Color online) The schematic diagram for a Brownian particle that walks in a discrete ratchet potential coupled with a linearly decreasing temperature heat bath. The temperature for the heat bath decreases from *T*_*h*_ to *T*_*c*_ according to Eq. (60).

For two ratchet potentials (*M* = 2), **P** has a form

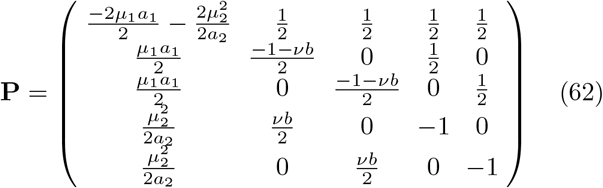

while for *M* = 3 and *M* = 4 cases, one gets

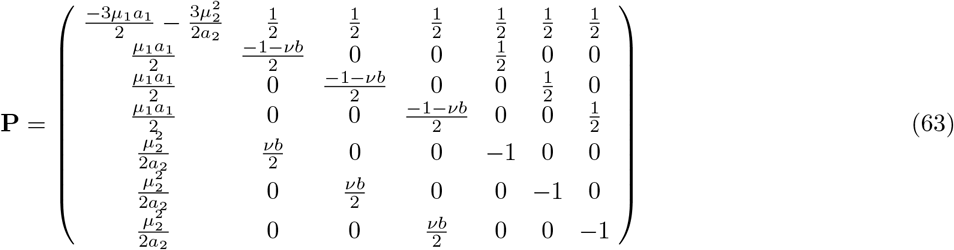

and

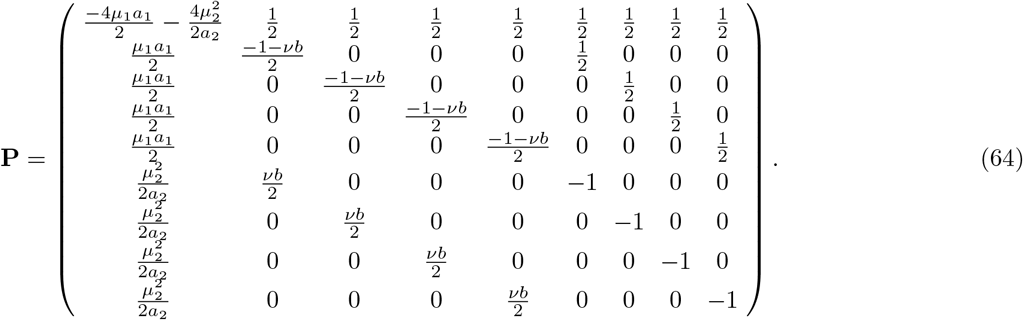

Here 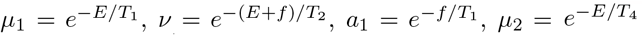 and 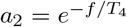. Since the temperature linearly decreases, the parameter *T*_1_ = *T*_*h*_, *T*_2_ = *T*_*h*_ +(*T*_*c*_ −*T*_*h*_)*/*3, *T*_3_ = *T*_*h*_ + 2(*T*_*c*_ − *T*_*h*_)*/*3 and *T*_4_ = *T*_*c*_. The sum of each column of the matrix **P** is zero, Σ_*m*_ **P**_*mn*_ = 0 which reveals that the total probability is conserved: 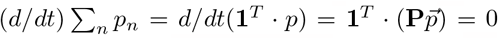. It is important to note that via the expressions *p*_1_(*t*), *p*_2_(*t*) and *p*_3_(*t*) that are shown in Appendix A2 and using the rates,

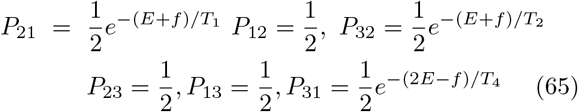

the thermodynamic quantities which are under investigation can be evaluated.

Once again, the velocity *V* (*t*) at any time *t* is the difference between the forward 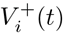 and backward 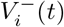 velocities at each site *i*

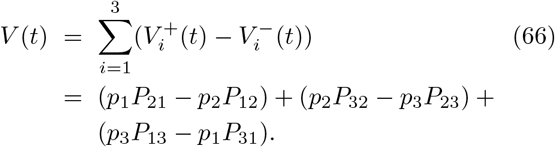

Similar to the previous section, as the number of network sizes increases, the entropy *S* of the system steps up. On the other hand, the velocity *V*, the entropy production *ė*_*p*_(*t*) as well as entropy extraction 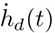 rates increase with network sizes. At steady state *V, ė*_*p*_(*t*) and 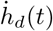 become independent of the network size. For all *M*, at steady the velocity converges to

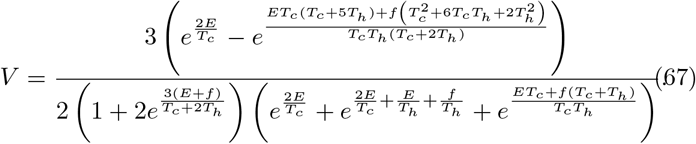

The energetics of the system that operates between the hot and cold baths are also compared and contrasted with a system that operates in a heat bath where its temperature linearly decreases along the reaction coordinate. We show that a system that operates between the hot and cold baths has significantly lower velocity but a higher efficiency in comparison with a linearly decreasing case as shown in Fig. 15. In the figure the ratio of velocity for a linearly decreasing case over the velocity for two heat baths cases is plotted as a function of *t* and *τ* for a given *ϵ* = 2.0, *M* = 4 and *f* = 0.3. The entropy for the linearly decreasing case is considerably larger than the entropy for two heat bath cases as shown in Fig. 16.

**FIG. 15:**
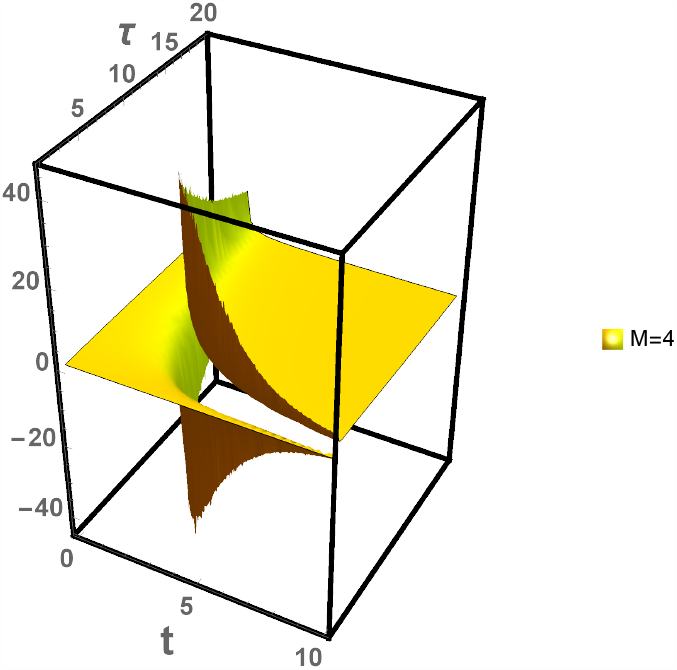
(Color online) The ratio of velocity for linearly decreasing case over the velocity for two heat baths case is plotted as a function of *t* and *τ* for a given *ϵ* = 2.0 and *f* = 0.3. In the figure, *M* is fixed as *M* = 4.

**FIG. 16:**
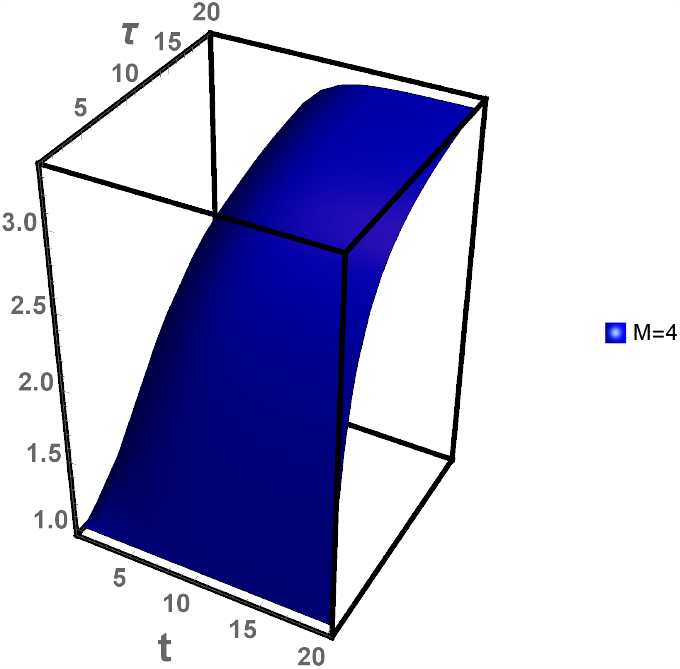
(Color online) The ratio of entropy for linearly decreasing case over the velocity for two heat baths case as a function of *t* and *τ* for a given *ϵ* = 2.0 and *f* = 0.3. In the figure, *M* is fixed as *M* = 4.

As depicted in Fig. 17a, a system that operates between the hot and cold baths has a significantly lower entropy production rate than a linearly decreasing temperature case. Moreover, in Fig. 17b, we compare the ratio of the free energy rate of a linearly decreasing case with a system that operates in two heat bath cases. The figure depicts that the free energy rate is considerably large for a linearly decreasing case. Our analysis also indicates that the change in free energy becomes minimal at a steady state. However, for a system that operates in a heat bath where its temperature varies linearly in space, the change in free energy varies linearly in space. The fact that the entropy, entropy production, and extraction rates are considerably larger for a a linearly decreasing temperature case than a Brownian particle that operates between the hot and cold baths indicates that systems exposed to a linearly decreasing temperature case are inherently irreversible and as a result, such systems have very low efficiency. On the contrary, the particle moving in a linearly decreasing thermal arrangement has a higher velocity. Thus such kind of thermal arrangement is beneficial, to designing a device that can transport a Brownian particle fast along the reaction coordinate. A Brownian particle that operates between two heat baths has a higher efficiency but a lower velocity in comparison with a linearly decreasing temperature case. Thus such a model system is advantageous if the sole purpose is to design an efficient Brownian motor.

**FIG. 17:**
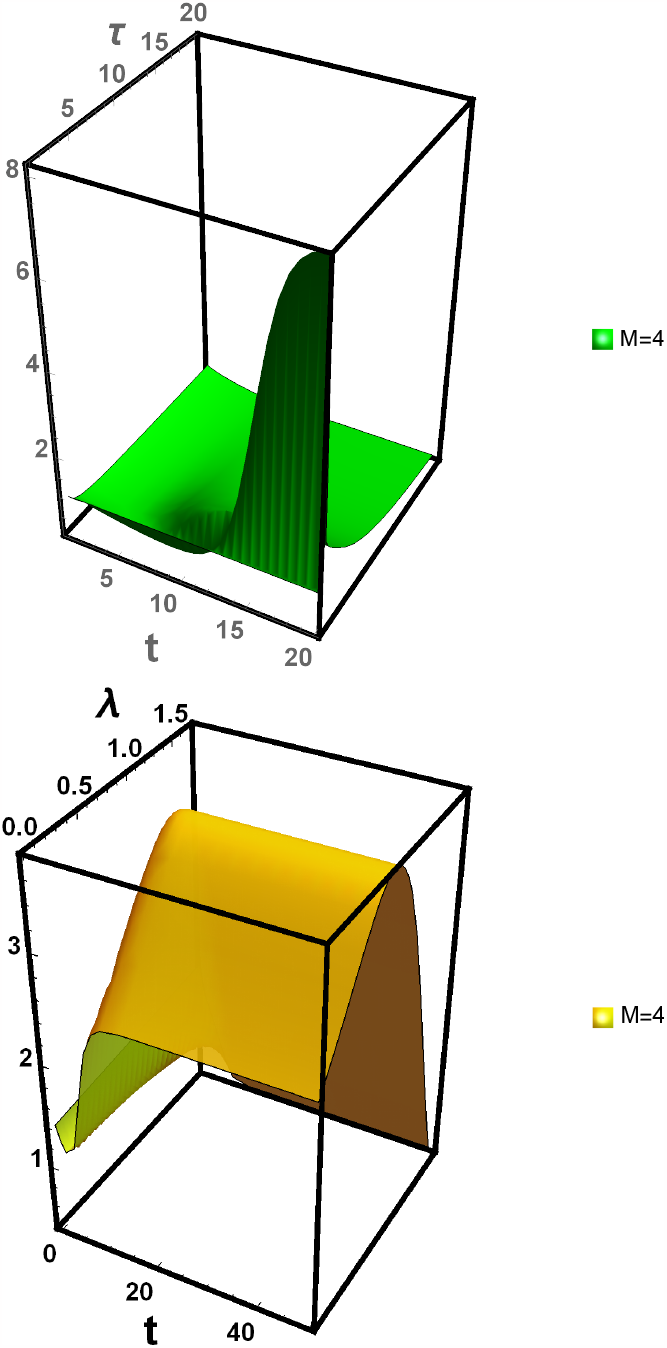
(Color online) (a)The ratio of entropy production rate for linearly decreasing case over the entropy production rate for two heat baths case as a function of *t* and *τ*. (b) The ratio of free energy rate for linearly decreasing case over the free energy rate for two heat baths cases as a function of *t* and *τ*. In the figures, the parameters are fixed as *M* = 4, *ϵ* = 2.0, and *f* = 0.3.

## IV. SUMMARY AND CONCLUSION

Most of the previous works have focused on exploring the thermodynamics feature of systems that operate in a single thermal ratchet. On the contrary, molecular motors such as kinesin, myosin, and dynein walk in microtubule networks. Most Biologically important problems such as impurities (donors) diffuse along the semiconductor layer and consequently, they require 2*D* and 3*D* analysis. Nanorobots are artificial Brownian motors and they are designed to achieve a unidirectional motion on complex networks. To address such important physical problems, an exactly solvable model is presented. The long-time, as well as the short-time behavior of the system, is explored by obtaining exact time-dependent solutions.

To comprehend the thermodynamic features of systems beyond a linear response and steady-state the regime, the thermodynamic features of a single Brownian particle that hops along M Brownian ratchets arranged in complex networks are studied. Each ratchet potential is either coupled with hot and cold reservoirs or a heat bath where its temperature decreases linearly along the reaction coordinate. Often the thermodynamic quantities such as entropy *S*(*t*), entropy production *e*_*p*_(*t*), and entropy extraction *h*_*d*_(*t*) serve as an indicator of irreversibly that the system sustains. Whenever our system obeys a detail balance condition, *e*_*p*_ = 0. On the contrary, when the system operates at a finite time or when it is exposed to symmetry-breaking fields, the entropy of systems can be greater than zero *e*_*p*_ *>* 0. Our analysis indicates that *S, e*_*p*_(*t*) and entropy extraction *h*_*d*_(*t*) step up as the network size *M* increases. On the contrary, the rates for thermodynamic relations such as velocity, entropy production rate as well as entropy extraction rate becomes independent of the network size as long as a periodic the boundary condition is imposed.

Furthermore, the thermodynamic properties of systems that operate between hot and cold baths are also compared with a system that operates in a heat bath where its temperature linearly decreases along with the reaction coordinate. Our result depicts that, regardless of the network size M the magnitude of the thermodynamic quantities such as entropy production and entropy extraction is also considerably larger for linearly decreasing temperature case than a system that operates between the hot and cold baths. This reveals that the degree of irreversibility is higher for the system that operates in a heat bath where its temperature decreases linearly along with the reaction coordinate.

The exactly solvable model presented in this paper serves as a tool to explore the thermodynamic features of two-dimensional systems. When the Brownian particle is exposed to a network coupled with two heat baths, the particle (motor) exhibits a higher efficiency but it achieves its task steadily. When the particle is exposed to a heat bath where its temperature decreases linearly, it moves faster but has lower efficiency. For all thermal arrangements, the particle attains a faster velocity whenever the network M increases. This indicates that by properly arranging the model ingredient prior to the motor operation, the motor can accomplish is a specific task. Thus the exactly solvable model presented in this work not only advances the nonequilibrium statistical mechanics but also helps in designing artificial Brownian motors.

## Appendix A1

In this Appendix we will give the expressions for *p*_1_(*t*), *p*_2_(*t*) and *p*_3_(*t*) as well as *V* (*t*) for a Brownian particle that operates between the hot and cold baths. For the particle which is initially situated at site *i* = 2, the time dependent normalized probability distributions after solving the rate equation 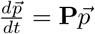 are calculated as

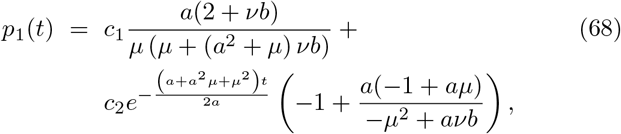

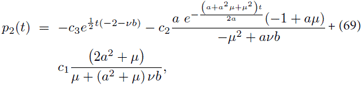

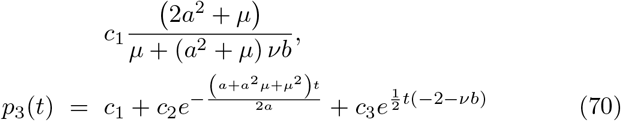

where

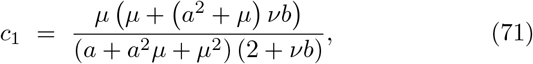

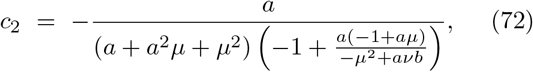

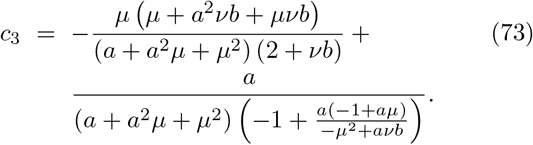

Here 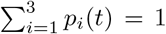 revealing the probability distribution is normalized. In the limit of *t* → ∞, we recapture the steady state probability distributions

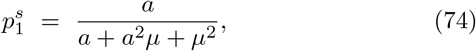

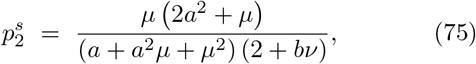

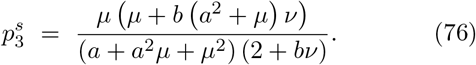

The velocity *V* (*t*) at any time *t* is the difference between the forward 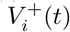 and backward 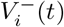 velocities at each site *i*

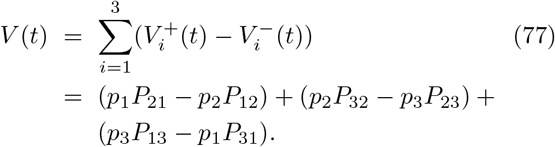

Exploiting Eq. (63), one can see that the particle attains a unidirectional current when *f* = 0 and *T*_*h*_ *> T*_*c*_. For isothermal case *T*_*h*_ = *T*_*c*_, the system sustains a non-zero velocity in the presence of load *f* ≠ 0 as expected. Moreover, when *t* → ∞, the velocity *V* (*t*) increases with *t* and approaches the steady state velocity

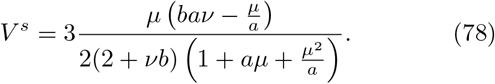

## Appendix A2

The expressions for *p*_1_(*t*), *p*_2_(*t*) and *p*_3_(*t*) as well as *V* (*t*) are derived considering a Brownian particle that operates in a heat bath where its temperature decreases linearly along with the reaction coordinate. For the particle which is initially situated at site *i* = 2, the time-dependent normalized probability distributions after solving the rate equation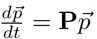 are given as

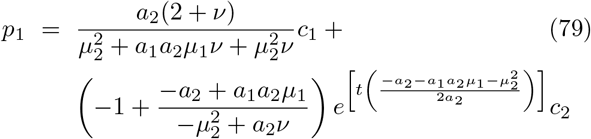

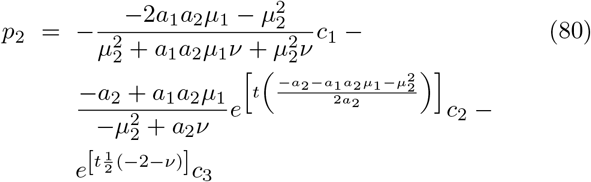

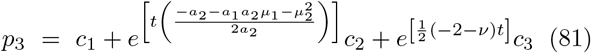

where

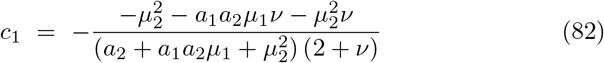

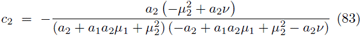

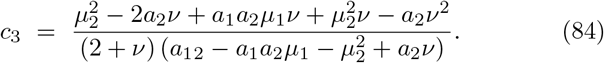

Once again, 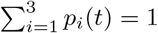 revealing the probability distribution is normalized. When *t*→ ∞, the steady state probability distributions converge to

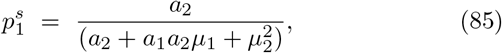

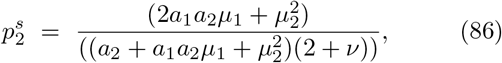

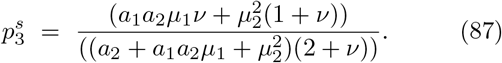

The velocity *V* (*t*) at any time *t* is the difference between the forward 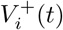 and backward 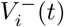 velocities at each site *i*

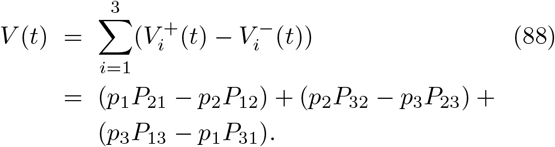

In the limit *t*→ ∞, the velocity *V* (*t*) increases with *t* and approaches to steady state velocity

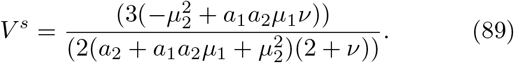

## Appendix A3

The expressions for *p*_1_(*t*), *p*_2_(*t*), *p*_3_(*t*), *p*_4_(*t*) and *p*_5_(*t*) as well as *V* (*t*) are derived considering a Brownian particle that operates in a cold and hot heat baths along with two dimensional lattice. For the particle which is initially situated at site *i* = 2, the time-dependent normalized probability distributions after solving the rate equation 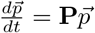are given as

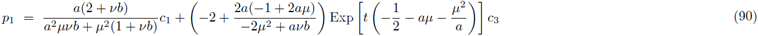

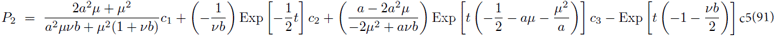

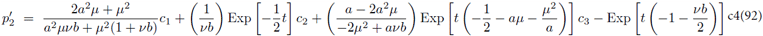

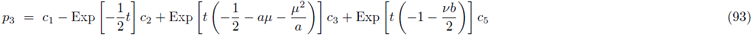

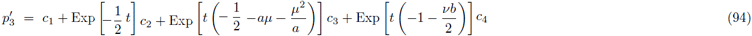

where

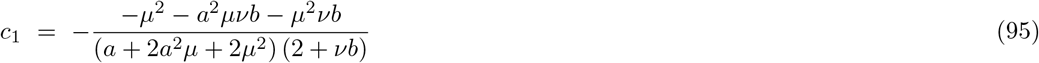

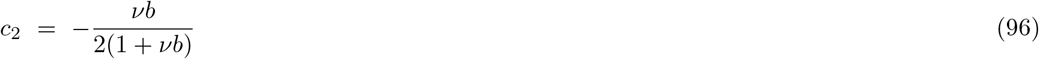

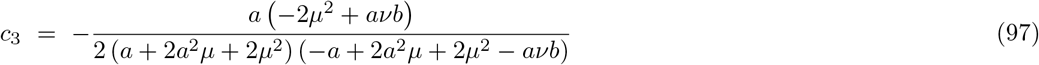

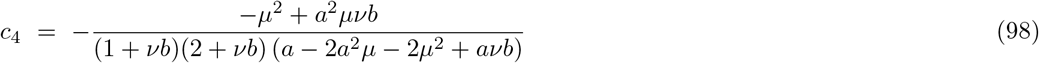

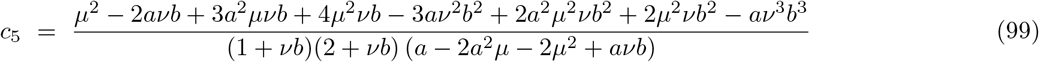

Once again 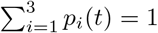 revealing the probability distribution is normalized. When *t* → ∞, the steady state probability distributions converge to

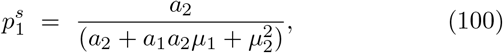

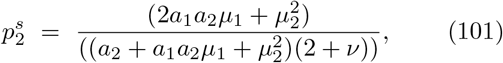

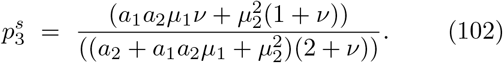

The velocity *V* (*t*) at any time *t* is the difference between the forward 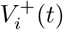 and backward 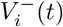velocities at each site *i*

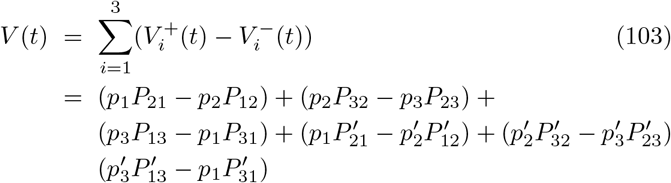

In the limit *t* → ∞, the velocity *V* (*t*) increases with *t* and approaches to steady state velocity

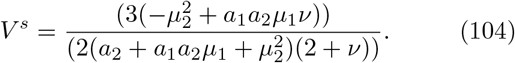

## Appendix A4

The expressions for *p*_1_(*t*), *p*_2_(*t*), *p*_3_(*t*), *p*_4_(*t*) and *p*_5_(*t*) as well as *V* (*t*) are derived considering a Brownian particle that operates in a linearly decreasing heat baths along with two dimensional lattice. For the particle which is initially situated at site *i* = 2, the time-dependent normalized probability distributions after solving the rate equation 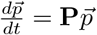 are given as

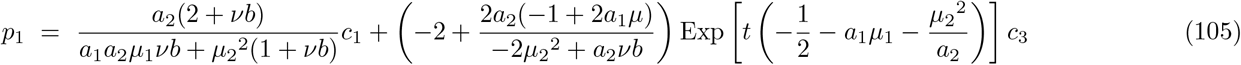

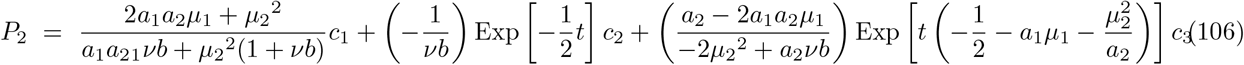

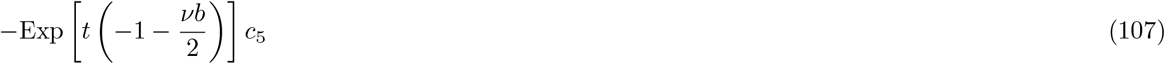

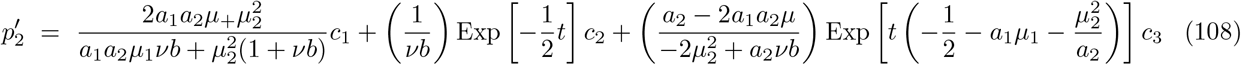

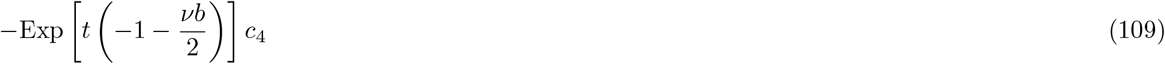

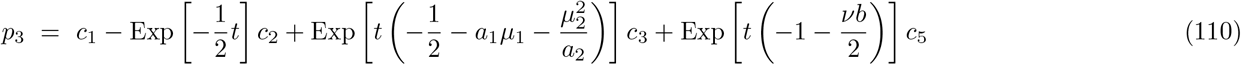

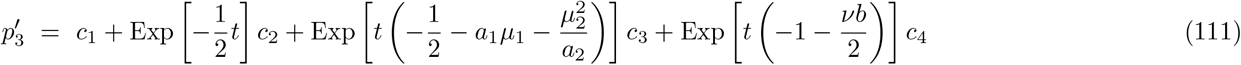

where

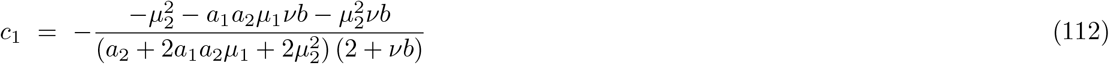

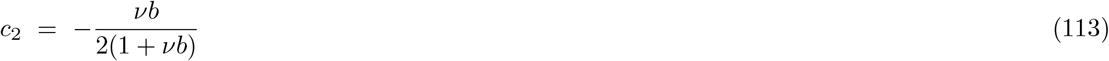

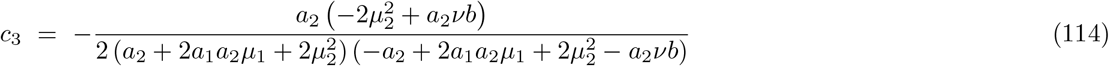

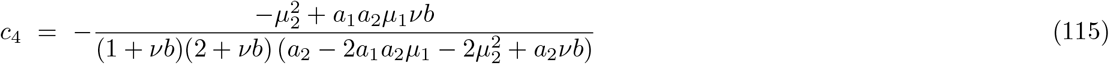

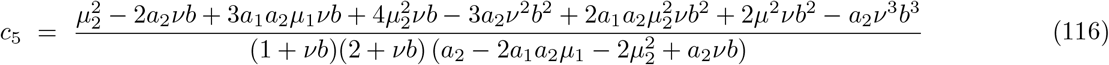

Once again,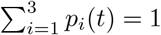 revealing the probability distribution is normalized. When *t* → ∞, the steady state probability distributions converge to

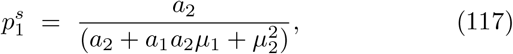

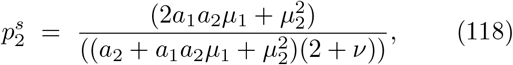

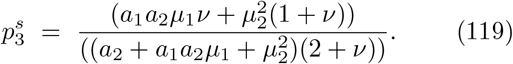

The velocity *V* (*t*) at any time *t* is the difference between the forward 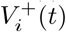 and backward 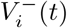 velocities at eachsite *i*

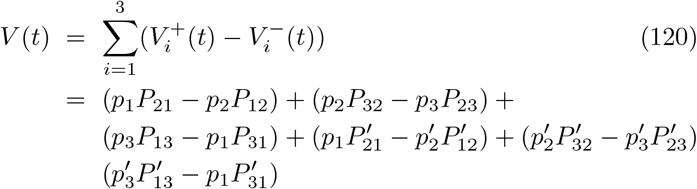

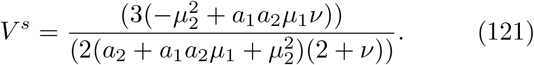

## Acknowledgment

I would like to thank Hana Taye and Mulu Zebene for their constant encouragement.

## References

[1] R. Clausius, Annalen der Physik 201, 353 (1865).

[2] G. T. Landi and M. Paternostro, Rev. Mod. Phys. 93, 035008 (2021).

[3] L. Onsager, Phys. Rev.37, 405 (1931).

[4] L. Onsager, Phys. Rev. 38, 2265 (1931).

[5] H. Ge and H. Qian, Phys. Rev. E 81, 051133 (2010).

[6] T. Tome and M. J. de Oliveira, Phys. Rev. Lett. 108, 020601 (2012).

[7] J. Schnakenberg, Rev. Mod. Phys. 48, 571 (1976).

[8] T. Tome and M.J. de Oliveira, Phys. Rev. E 82, 021120 (2010).

[9] R.K.P. Zia and B. Schmittmann, J. Stat. Mech. P07012 (2007).

[10] U. Seifert, Phys. Rev. Lett. 95, 040602 (2005).

[11] T. Tome, Braz. J. Phys. 36, 1285 (2006).

[12] G. Szabo, T. Tome and I. Borsos, Phys. Rev. E 82, 011105 (2010).

[13] B. Gaveau, M. Moreau and L.S. Schulman, Phys. Rev. E 79, 010102 (2009).

[14] J.L. Lebowitz and H. Spohn, J. Stat. Phys. 95, 333 (1999).

[15] D. Andrieux and P. Gaspar, J. Stat. Phys. 127, 107 (2007).

[16] R.J. Harris and G.M. Schutz, J. Stat. Mech. P07020 (2007).

[17] Tania Tome and Mario J. de Oliveira, Phys. Rev. E 9, 042140 (2015).

[18] J.-L. Luo, C. Van den Broeck, and G. Nicolis, Z. Phys. B 56, 165 (1984).

[19] C.Y. Mou, J.-L. Luo, and G. Nicolis, J. Chem. Phys. 84, 7011 (1986).

[20] C. Maes and K. Netocny, J. Stat. Phys. 110, 269 (2003).

[21] L. Crochik and T. Tome, Phys. Rev. E 72, 057103 (2005).

[22] M. Asfaw, Phys. Rev. E 89, 012143 (2014).

[23] M. Asfaw, Phys. Rev. E 92, 032126 (2015).

[24] K. Brandner, M. Bauer, M. Schmid and U. Seifert, New. J. Phys. 17, 065006 (2015).

[25] B. Gaveau, M. Moreau and L. S. Schulman, Phys. Rev. E 82, 051109 (2010).

[26] E. Boukobza and D.J. Tannor, Phys. Rev. Lett. 98, 240601 (2007).

[27] M. A. Taye, Phys. Rev. E 94, 032111 (2016).

[28] M. A. Taye, Phys. Rev. E 101, 012131 (2020).

[29] H. Ge, Phys. Rev. E 89, 022127 (2014).

[30] H. K. Lee, C. Kwon, and H. Park, Phys. Rev. Lett. 110, 050602 (2013).

[31] R. E. Spinney and I. J. Ford, Phys. Rev. Lett. 108,170603 (2012).

[32] H. K. Lee, C. Kwon and H. Park, Phys. Rev. Lett. 110, 050602 (2013).

[33] A. Celani, et al., Phys. Rev. Lett. 109, 260603 (2012).

[34] P. Strasberg and M. Esposito, Phys. Rev. E 99, 012120 (2019).

[35] S. K. Manikandan, D. Gupta, and S. Krishnamurthy, Phys. Rev. Lett. 124, 120603 (2020).

[36] D. J. Skinner and J. DunkelPhys. Rev. Lett. 127, 198101 (2021).

[37] S. Otsubo, S. Ito, A. Dechant, and T. Sagawa, Phys. Rev. E 101, 062106 (2020).

[38] T. V. Vu, V. T. Vo, and Y. Hasegawa, Phys. Rev. E 101, 042138 (2020).

[39] T. Koyuk and U. Seifert, Phys. Rev. Lett. 122, 230601 (2019).

[40] M. A. Taye, Phys. Rev. E 105, 054126 (2022).

[41] T. Bameta, D. Das, R. Padinhateeri and M. M. Inamdar, ArXiv:1503.06529 (2015).

[42] D. Oriola and J. Casademunt, Phys. Rev. Lett. 111, 048103 (2013).

[43] O. Campa, Y. Kafri, K.B. Zeldovich, J. Casademunt and J.-F. Joanny, Phys. Rev. Lett. 97, 038101 (2006).

[44] H. Ge, Phys. Rev. E 89, 022127 (2014).

[45] I. Goldhirsch and Y. Gefen, Phys. Rev. A 33, 2583 (1986).

[46] I. Goldhirsch and Y. Gefen, Phys. Rev. A 35, 1317 (1987).

[47] M. A. Taye, Eur. Phys. J. B 94, 124 (2021).

[48] M. A. Taye, J. Stat. Phys. 169, 423 (2017).

